# Trapping non-cognate nucleotide upon initial binding for replication fidelity control in SARS-CoV-2 RNA dependent RNA polymerase

**DOI:** 10.1101/2023.09.01.555996

**Authors:** Moises E. Romero, Shannon J. McElhenney, Jin Yu

## Abstract

The RNA dependent RNA polymerase (RdRp) in SARS-CoV-2 is a highly conserved enzyme responsible for viral genome replication/transcription. Here we investigate computationally natural non-cognate vs cognate nucleotide addition cycle (NAC) and intrinsic nucleotide selectivity during the viral RdRp elongation, focusing *prechemically* from initial nucleotide substrate binding (enzyme active site open) to insertion (active site closed) of RdRp in contrast with one-step only substrate binding process. Current studies have been first carried out using microsecond ensemble equilibrium all-atom molecular dynamics (MD) simulations. Due to slow conformational changes (from the open to closed) accompanying nucleotide insertion and selection, enhanced or umbrella sampling methods have been further employed to calculate free energy profiles of the non-cognate NTP insertion. Our studies show notable stability of noncognate dATP and GTP upon initial binding in the active-site open state. The results indicate that while natural cognate ATP and Remdesivir drug analogue (RDV-TP) are biased to be stabilized in the closed or insertion state, the natural non-cognate dATP and GTP can be well trapped in *off-path* initial binding configurations. Current work thus presents an intrinsic nucleotide selectivity mechanism of SARS-CoV-2 RdRp for natural substrate fidelity control in viral genome replication.

## 1 Introduction

The SARS-CoV-2 virus responsible for the COVID-19 pandemic continues to evolve [1] and pose a threat to human life [2]. While the vaccines developed are demonstrating success, there remains an imperative to accelerate antiviral development on therapeutics, considering virus’s evolution and resistance, vaccine hesitancy among individuals, or the inability of some countries to afford the available vaccines. While much of the drug development is focused on targeting the viral spike protein [3–5] or the main protease [6–8], there are significant challenges. The spike protein is known for its high variability [1, 9, 10] and the protease [11] can also mutate, becoming resistant to drugs, as seen in HCV [12, 13]. In contrast, the core replication machinery of SARS-CoV-2, the RNA dependent RNA polymerase (RdRp) or nonstructural protein nsp12, is a highly conserved drug target [14, 15]. Here we focus on studying SARS-CoV-2 RdRp to understand its underlying function and mechanism that is crucial for future drug development [16], particularly given the significant gaps in knowledge regarding the functioning of viral RdRps [17–19].

Upon the pandemic upheaval in 2020, a few high-resolution cryo-EM structures of SARS-CoV-2 RdRp were released immediately, including apo with the RdRp active site open [20,21] and a post-catalysis form with the active site closed [21]. The post-catalysis structure was bound with a nucleotide drug analogue from remdesivir (RDV) [21]. These structures were in complex with segments of the nsp7/nsp8 cofactors. Later, additional high-resolution structures were resolved with longer nsp8 being “sliding poles” [22], as well as RdRp in conjunction with the nsp13 helicase enzyme [23], and in both pre- and post-translocation states [24]. Further structures illustrated RdRp backtracking [25] or stalling [26] due to the interaction with the drug analogue RDV. Similarly, there were structures obtained with favipiravir, another nucleotide analogue drug [27, 28]. Overall, these structures adopt the post-catalysis state [21] in the nucleotide addition cycle (NAC) of the viral RdRp, leaving the initial nucleotide binding (active site open) and pre-catalytic insertion or substrate (closed) states unresolved. It was until very recently that the insertion state with ATP as a cognate nucleotide triphosphate (NTP) bound in the active site was resolved [29]. Currently, the initial binding or open state structure of RdRp remains unresolved, although such a structure has been identified for the Poliovirus [30] (PV) or Enterovirus [31] (EV) RdRp, which share structural similarities with SARS-CoV-2 RdRp.

Efforts to identify drug inhibitors for the SARS-CoV-2 RdRp have been made extensively [16, 32, 33]. Computational docking has often been employed, which predominantly focus on nucleotide immediate binding to an apo-form RdRp structure [34,35], non-differentiating the active open or closed form, and overlooking initially the functional RdRp (nsp12) elongation complex composed additionally of nsp8, nsp7, and RNA strands. Moreover, while atomic molecular dynamics (MD) studies have provided insights into interactions of nucleotide analogues with the viral RdRp, they often utilize directly the insertion state [36–38], ignoring the initial nucleotide analogue binding stage that can be essential for nucleotide screening or selectivity upon entry. Meanwhile, single-molecule studies have offered a glimpse into the dynamics of the elongation cycle, revealing that RdRp can adopt fast, slow, or very slow catalytic pathways with variable rates contingent upon the kinetic pathway [39]. Additionally, information in regard to the RdRp translocation in the NAC has advanced through the cryo-EM studies, which demonstrated a structural rearrangement in nsp8 to accommodate the exiting RNA duplex [24]. Computational work has shown that the incorporation of RDV-TP into the primer strand results in a steric clash at the conserved motif-B of the RdRp leading to an unstable post-translocation state int comparison with pre-translocation, i.e., as a mechanism for antiviral analogue termination of elongation [40]. An alternative suggestion based on single-molecule studies [39] proposes that RdRp backtracks up to 30 nucleotides (nts) after RDV-TP incorporation, which can be interpreted as elongation termination in standard assays. Despite these efforts on quantitative studies of the RdRp NAC, a critical gap remains in understanding initial nucleotide substrate binding to insertion, which is fundamental for nucleotide selectivity and antiviral drug design, given the substrate screening and pre-chemistry inhibition as essential fidelity checkpoints in stepwise NAC [41–43].

Accompanied with the nucleotide substrate binding to insertion, a substantial protein conformational transition happens, which is likely to be a rate limiting step in the NAC, as demonstrated in structurally similar single-subunit RNA or DNA polymerases (RNAPs or DNAPs) [44–46]. Such a rate-limiting pre-chemical step accordingly plays a highly essential role in the nucleotide substrate selectivity [47]. To quantify the process with energetics, we calculated the free energy profile or the potential of mean force (PMF) of the nucleotide insertion, recently for cognate substrate ATP and the corresponding nucleotide drug analogue RDV-TP [48]. We found that both ATP and RDV-TP become significantly more stabilized in the insertion state than upon initial binding. In contrast to natural substrate ATP, RDV-TP behaves differently in its interaction with the template nt. Cognate ATP forms the Watson-Crick (WC) base pairing with template uracil at the +1 position, from the initial binding through to insertion. On the other hand, RDV-TP initially forms base stacking with the template nt at +1 upon binding. It then inserts into the active site, facing an energetic barrier that is marginally low or comparable to that of cognate ATP [48].

Note that RDV-TP analogue differs from the cognate ATP by only a few atoms: with a 1’ cyano group attached on the sugar C1’ and 3 atomic replacements on the base. Interestingly, we have also noticed that one conserved motif-F (R553+R555) essentially facilitates the insertion of cognate ATP via interactions with the ATP-triphosphate tail. While such phosphate interactions could potentially hinder the drug analogue RDV-TP insertion, it was subtly avoided upon sufficient thermal fluctuations from the template nt+1 that forms base stacking with RDV-TP [48].

In current work, we focus on characterizing the intrinsic nucleotide polymerase enzyme selectivity of SARS-CoV-2 RdRp, i.e., the selectivity or differentiation between natural cognate NTP (ATP here) and natural non-cognate NTP substrates. To do that, we examined the nucleotide insertion dynamics of non-cognate dATP and GTP, in comparison with cognate ATP and RDV-TP analogue, and calculated the insertion PMFs of dATP and GTP starting from initial binding stage. Since the polymerase NAC lasts over tens of milliseconds in general [39], the rate-limiting transition accompanies the nucleotide binding to insertion (or active site from open to closed) of RdRp is expected to be on the millisecond timescale [44–46]. Therefore, such an insertion process cannot be sampled directly using equilibrium MD simulation that is limited by the sub-microsecond to microseconds timescale [49, 50]. To obtain free energetics of such a dynamics process, we extended our previous methodology on employing umbrella sampling MD simulations to construct the PMFs of various NTPs from initial binding to insertion [48,51]. We first constructed atomic structural models of the RdRp-nsp7-nsp8-RNA complexes bound with the noncognate nucleotide (ncNTP) species, in both initial binding and insertion states. Subsequently, we performed all-atom equilibrium simulations at sub-microseconds in ensembles to characterize the respective initial binding and insertion states of the non-cognate dATP/GTP bound RdRp complexes. Lastly, we obtained the PMFs of the dATP/GTP insertion processes using the umbrella sampling MD simulations, following initial insertion paths constructed on top of collective reaction coordinates (RCs), according to displacements of atomic coordinates from seven highly conserved structural motifs of RdRp (A to G) and incoming NTP (with and without the associating template nt). Our aim is to elucidate the intrinsic nucleotide selectivity of SARS-CoV-2 RdRp, which turns out to be primarily relying on ‘trapping’ the non-cognate nucleotide species upon entry or initial binding to certain configurations at the active site and preventing them from further insertion.

## 2 Methods

### 2.1 Computational Details

#### 2.1.1 Constructing Open and Closed models for dATP and GTP

Based on the approach used in previous work on modeling SARS-CoV-2 RdRp [48], we constructed models for initial binding (open) and insertion (closed) of various NTPs using high-resolution Cryo-EM structures of SARS-CoV-2 nsp12-nsp7-ns8-RNA complexes. The model for RDV-TP in the insertion state was created using a post-catalytic structure bound with RDV-TP (PDBid:7BV2) as a reference, which also includes catalytic Mg^2+^ ions [21]. For the ATP insertion model, we positioned ATP over the RDV-TP in the insertion model. The RDV-TP initial binding model was created using an apo structure (PDBid:7BTF) [20] aligned with the RDV-TP insertion structure, with RNA, RDV-TP, and Mg^2+^ ions copied over. The ATP initial binding model was built following the same process as described above. The dATP states were built using the ATP models as reference and removing the 2’ OH group. Alternatively, GTP was aligned with the ATP models with subsequent geometry optimization to force wobble (WB) base pairing (see **Figure 2** *lower right*) [52]. For complete details on preparation of RdRp model in association with RDV-TP (with force-field parameterization) and ATP, please see previous work [48, 53].

#### 2.1.2 Simulation Parameters and Setup

All MD simulations were performed using GPU-accelerated Gromacs 2021 software [54] with the following forcefields: Amber14sb [55], Parmbsc1 [56], triphosphate parameters developed previously [57]. Enhanced or umbrella sampling methods were performed using a Gromacs 2021 package patched with PLUMED [58]. Each complex was solvated with explicit TIP3P water [59] in a cubic box with a minimum distance from complex to the wall of 15 Å. Resulting in an average box dimensions of 15.7nm x 15.7 nm x 15.7 nm. The overall negative charge of the complex was neutralized and ions were added to create a salt concentration environment of 100mM. Full simulation systems were *∼*382,000 atoms in size. A cut-off of 10Å was used to treat short range electrostatics interactions and the Particle-mesh-Ewald (PME) algorithm to treat long range interactions [60]. The LINCS algorithm is used to constrain bonds to hydrogen atoms allowing the use of a 2 fs timestep when integrating the equations of motion [54]. Temperature was kept at 310K using the velocity rescaling thermostat. Pressure was kept at 1 bar using Berendsen barostat [61] during equilibration and Parrinello-Rahman barostat [62] for production, targeted MD (TMD), and umbrella simulation runs. Each system was minimized for a maximum of 50000 steps using steepest-descent algorithm, followed by a slow equilibration with restraints released (every 1 ns) going from NVT (canonical or constant volume and temperature) ensemble to NTP (constant pressure and temperature) as previously used [48].

For each NTP initial binding and inserted states ten 100 ns equilibration trajectories were launched, with 10 ns removed from the start, to create 900 ns for each NTP state, totaling *∼*7.2 *µ*s of simulation time for RDV-TP/ATP/dATP/GTP in both open and closed conformation states. The generated equilibrium ensemble was then used for analysis and selection of references for free energy calculations.

### 2.2 Reaction coordinate (RC), launching NTP insertion path, and construction the Potential of Mean Force (PMF)

The reaction coordinate (or RC) used in umbrella sampling [63,64] is the difference in RMSD (eq. 1) of a current frame (coordinates X) with respect to two reference structures, one in the open (NTP initial binding) and the other in the closed (NTP insertion) state, respectively.

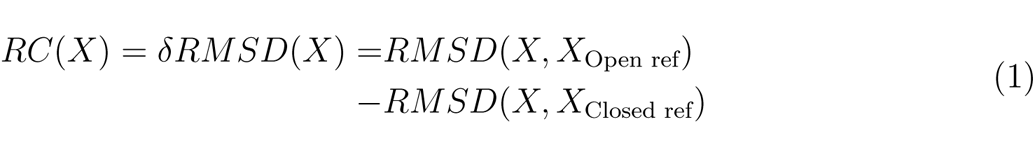

The reference structures (Open/Closed ref) were selected using the the first half (50 ns) of the equilibrium trajectory (or quasi-equilibrated). In the case of ncNTP we visually inspected to select a base pair geometry between the incoming NTP and template nt+1 from the well sampled or representative regions **Figure 2**.

After selecting the reference structures, a path of the NTP insertion is generated for umbrella sampling using target molecular dynamics (TMD) [65] simulations. We create a forward path (open to closed ref) and a backward path (closed to open), applying force along the following RC or atomic coordinates (*X*): nsp12 motifs (motif A-G backbone atoms), NTP (heavy atoms), and additionally tested two protocols [48]: i) include or with force on the template nt +1 ii) exclude or without force on the template nt +1.

From the TMD paths created between the two reference structures (with only the first half of the respective paths included to avoid large deviations from the references), intermediate structures were evenly (every 0.1 Å along the RC) selected to be used in launching umbrella sampling simulations. The force constants used in the TMD were carried over for umbrella sampling simulations (see **Table S2**). The collected RC value histograms were then re-weighted using the weighted histogram analysis method [66] as implemented by the WHAM package [67]. For each set of trajectories making up the umbrella windows, 10-ns trajectory was removed from the start, followed by convergence check for every 10ns. For dATP and GTP, the convergence time ranged from 50ns-60ns (**Figure S6**), in the case of no force applied on the template nt +1. GTP and dATP used 24 and 21 windows, respectively, for a total of *∼*1.2 *µ*s of simulation time per PMF. Additionally, error bars are estimated using the bootstrapping error analysis method [68] implemented in the WHAM package. For a thorough description on utilizing the umbrella sampling method for constructing the PMFs for ATP and RDV-TP insertion see previous work [48, 64].

To additionally examine the insertion structures simulated from umbrella sampling, we used steered molecular dynamics (SMD) to check whether the stability of the inserted GTP/dATP show consistencies with the umbrella sampling or the PMFs constructed. The SMD was implemented controlling two center of mass (COM) distances defined as between the heavy atoms of the NTP and the C*_α_* atoms from residues within 10 Å of the 3’ end primer. The distance between the two COMs was increased at a slow rate of 1 Å per 100 ns, with a force constant set to 2.4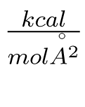.

### 2.3 Structural dynamics analyses

#### 2.3.1 RMSD

is measured for each RdRp complex on the subdomains (protein backbone), RNA (phosphate backbone), and NTP (heavy atoms) as on well as the the key motifs (from A to G) interacting with each NTP (see **Figures 1, S1, S2**). Both initial binding and insertion equilibrium ensemble trajectories were aligned via the finger’s subdomain to their respective initial states after minimization for each NTP.

**Figure 1:**
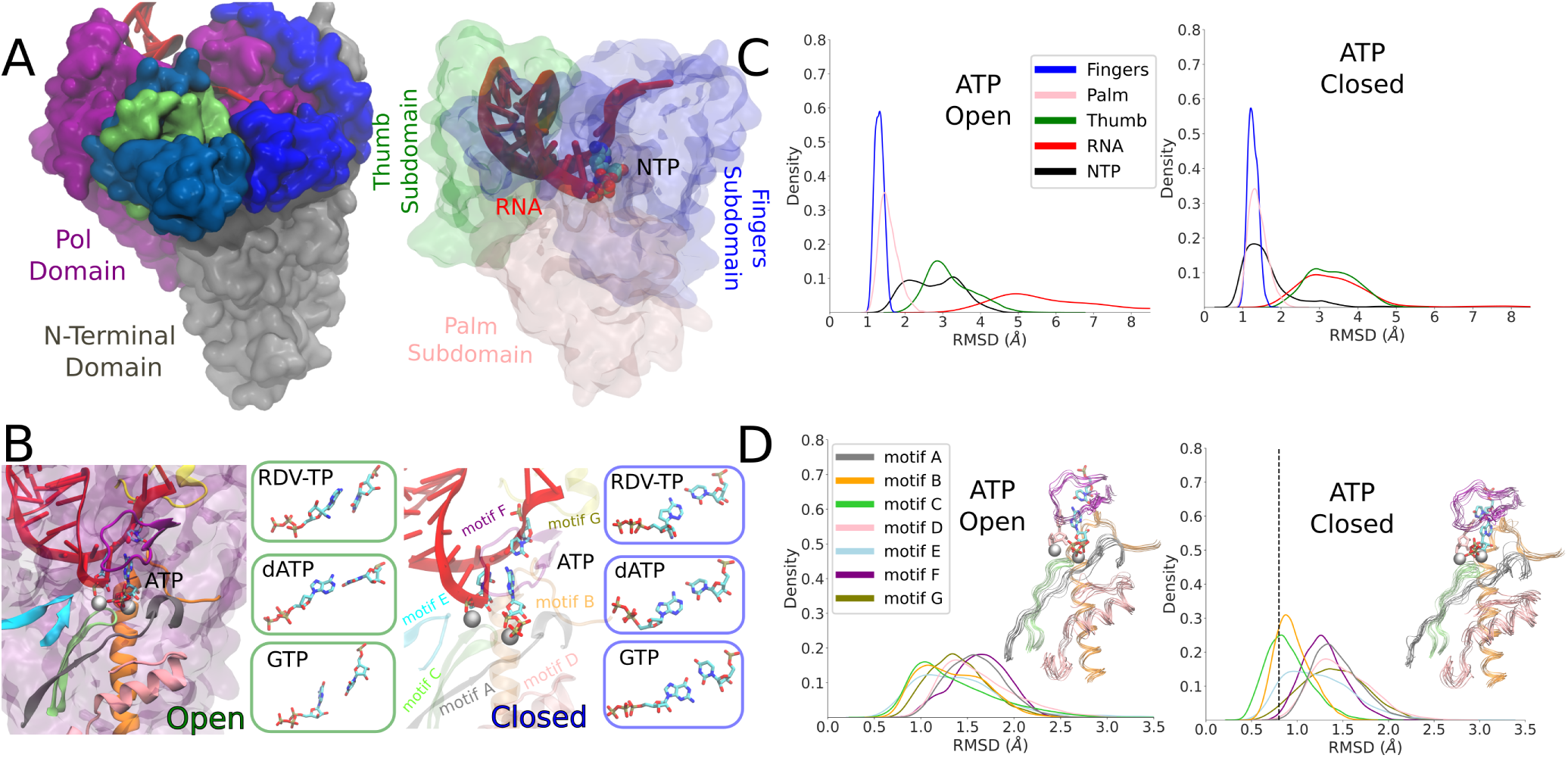
The constructed structural models of the SARS-CoV-2 RdRp elongation complex in its initial NTP substrate binding (open) and then inserted (closed) states. A: The simulated elongation complex is depicted with the nsp12 pol domain (purple) + N-terminal domain (gray), two cofactor nsp8 (blue) and nsp7 (green), RNA (red) and an NTP in the active site (*left*). The three major subdomains within the pol domain: fingers in blue, palm in pink, and thumb in green. The RNA is shown in red as well as a NTP shown in space filling spheres (*right*). B: The modeled and simulation equilibrated ATP is shown as bound initially to an open active site, with the seven protein motifs highlighted in color. The boxes to the right show the initially bound RDV-TP, dATP, and GTP that were also modeled and equilibrated from the simulations (*left*). The modeled and equilibrated ATP is shown in the insertion or the active site closed state, with the seven structures motifs shown as well, and the boxes to the right displaying the modeled and equilibrated insertion configurations of RDV-TP, dATP, and GTP to the closed active site (*right*). C: The subdomain RMSD for simulating initial binding (*left*) and insertion (*right*) for ATP, see **Figure S1** for the rest of the NTPs. D: The motif RMSD for initial binding (*left*) and insertion (*right*) for ATP, see **Figure S2** & **Table S1/S2** for the rest of the NTPs.

#### 2.3.2 Base Pair Geometry

was measured between the incoming NTP and uracil template nt at +1 by calculating the base plane angle (C1’-C7-C5 for NTP and C1’-C2-C5 for template) and the distance between the center of mass (COM) of each base. A similar protocol was followed for measuring the geometry between each NTP and the 3’ end primer: measuring the COM between bases and the base plane angle for (C1’-C7-C5 for NTP and C1’-C2-C5 for 3’ end primer). Measurements were conducted using the MDAnalysis python package [69] and gromacs gangle module.

#### 2.3.3 Hydrogen Bond Occupancy

Hydrogen bonds (HB) are measured using the gromacs 2021 module. A distance cutoff of *≤*3.5 *Å* between donor-acceptor involved and hydrogen-donor-acceptor angle cutoff of *≤* 30° is used as the criterion to indicate a proper hydrogen bond. Unique HB with an occupancy greater than 20% (within a 900 ns combined equilibrium ensemble trajectory) are considered for analysis.

Plots are created using python packages seaborn [70] and matplotlib [71].

## 3 RESULTS

### 3.1 Equilibrium ensemble simulations: protein structural variations and distinctive dynamical responses to different NTPs upon initial binding and insertion

Upon modeling the initial binding (open) and insertion (closed) complexes for the four NTPs, we conducted equilibrium all-atom MD simulations of 10 x 100 ns for each system. We began by measuring and comparing the RMSDs of the protein subdomains, RNA and NTP’s using the respective energy minimized structures for insertion or initial binding as references (see **Figure 1** & **SI Figure S1**). For all four NTP binding, the fingers subdomain RMSD is centered around *∼*1.3 Å and the palm subdomain remains similarly aligned with fingers, especially in the insertion state. The thumb subdomain has a comparatively wide distribution of RMSD in all cases, indicating conformational flexibility. All NTP display less variability in the insertion state, along with the fingers/palm subdomain (**SI Figure S1**). The cognate ATP and RDV-TP analogue both exhibit lower RMSDs than non-cognate dATP or GTP in the insertion state. For initial binding, the RMSDs of GTP along with the RNA scaffold are much larger than that for the other NTP binding. Additionally we compared the RMSDs of the seven conserved motifs (see **Figure 1**, **SI Figure S2** and **Table S1/S2**). Both motif B and C show comparatively low RMSDs in the insertion state. Motif-C demonstrates notable stability among all motifs for all inserted NTPs, likely due to it hosting the key catalytic residues (S759/D760/D761). The RMSDs of other structural motifs displayed significant variations upon initial binding of different NTPs. In the insertion state, motifs F and G show more distinctions dynamically than rest of motifs for different NTP substrates.

Next, we examined the base association or pairing geometry between the initially binding/insertion NTP and template +1 nt (**Figure 2**). In the insertion equilibrium ensemble (active site closed), we observed that the NTP-template distance distribution predominately centers at *∼*6.5 *Å* and base plane angle *∼*30°, resulting in either the stable Watson-Crick (WC) base pairing (for ATP/RDV-TP/dATP-template) or wobble base pairing (WB) interactions (for GTP-template uracil). The probability of WC/WB base pairing is high for ATP (69%), RDV-TP (94%) and GTP (70%), low for dATP (47%). Indeed, dATP upon insertion displays more flexibility than other NTPs in association with the template nt (see **Figure 2** *upper right*). In the NTP initial binding ensemble (active site open), the NTP-template association geometries vary significantly. Upon ATP initial binding, as a significant amount of WC population (48%) is identified, a comparable amount of un-paired or weakly paired (single HB) ATP-template uracil configurations are also present. Upon initial binding of RDV-TP, three configurations have been identified, with either the WC base pairing (36%) or base stacking (38%) being stabilized, and (26%) unpaired [48]. For non-cognate NTPs, the initially bound dATP marginally forms WC base pairing (12%) with the template nt, while GTP upon initial binding cannot forms stabilized WB base pairing with the template uracil.

**Figure 2:**
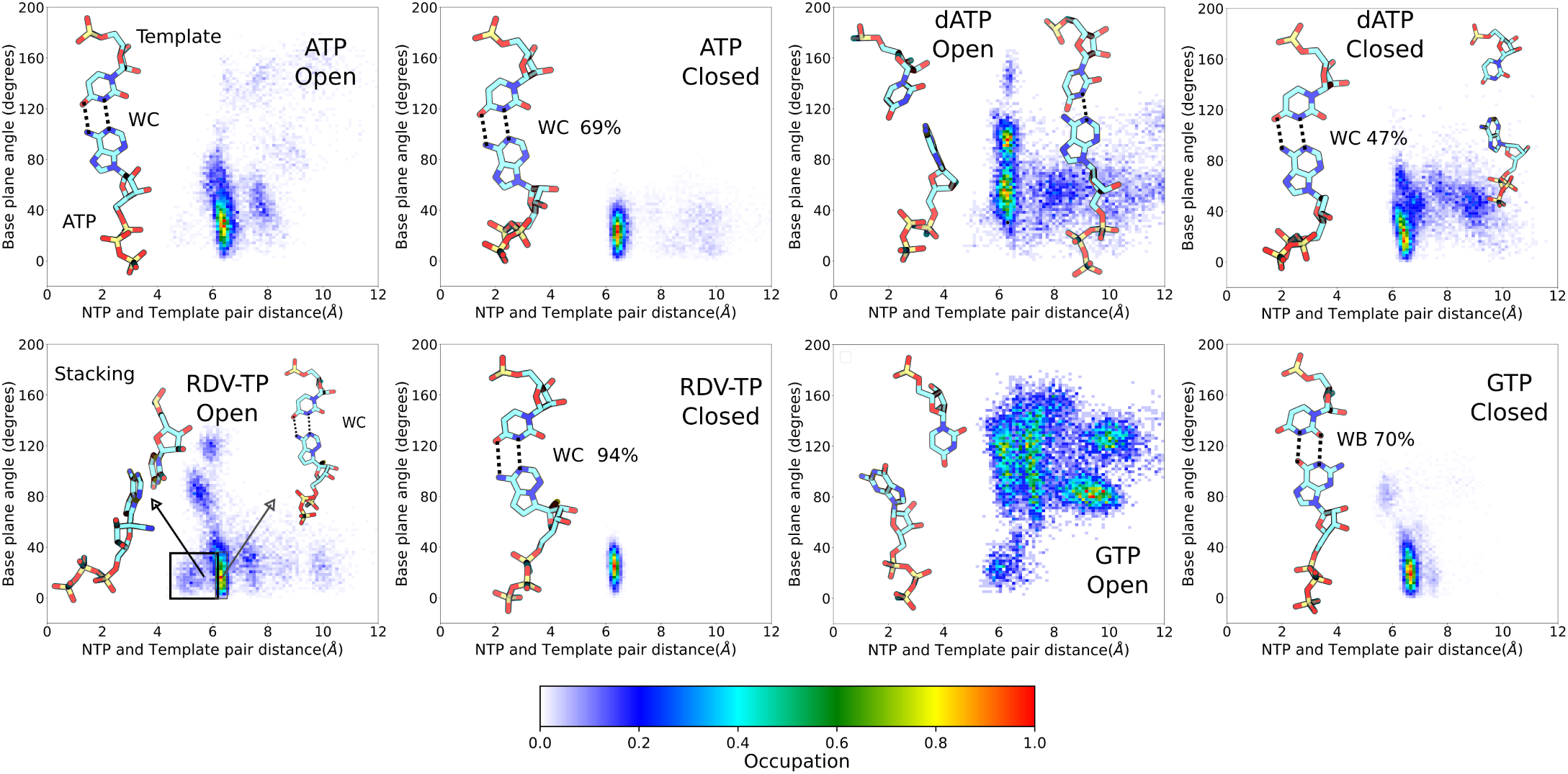
NTP-template association geometries from ensemble equilibrium simulations. The geometric measures (see **Methods**) between the uracil template +1 nucleotide (nt) and individual incoming NTP are displayed, for ATP (upper left), RDV-TP (lower left), dATP (upper right), and GTP (lower right), upon initial binding (left) and insertion (right) for each NTP species. Licorice representations of the NTP and template nt show the dominant geometries for each simulation system. Dotted lines indicate the hydrogen bonds for the standard Watson-Crick (WC) or wobble base (WB) pairing (percentile denoted).

Additionally, we measured the geometry between the NTP and 3’ end primer (**SI Figure S3**). From different NTP insertion ensembles, the NTP-3’ end primer distribution centers closely around *∼*4-6 *Å* and a base plane angle *∼*150° to 170°, showing stability. In contrast, upon initial binding, the mismatch GTP associated with the 3’ end primer in largely diverse geometries (distance spans from 4 to 13*Å* and the angle varies from 20° to 170°). The noncognate dATP upon initial binding also shows diverse geometries in association with the 3’ end primer (distance spans from 4 to 9 *Å* and the angle varies from 40° to 180°). For cognate ATP and RDV-TP, the initial binding geometries with respect to the 3’ end primer are much more localized, i.e., the distance dominantly spans from 4 to 5 *Å*, with small populations in between 8-10 *Å*; the angle varies from around 100° to 180° below than the non-cognate species. In summary, in the active site closed state or insertion equilibrium ensemble, various NTPs display much less variation of geometries with respect to the template +1 nt or the 3’ end primer than those in the active site open state or initial binding equilibrium ensemble. The highly diverse configurations of initially bound NTP (particularly non-cognate NTP) suggest that detection of various species of incoming NTP starts well from the beginning, i.e., upon the initial NTP binding when the active site remains open.

Furthermore, we also measured the HB occupancies for various NTPs in respective associations with the protein, template +1 nt, and 3’ end RNA primer (histogram statistics shown in **SI Figure S4**), for both the NTP initial binding and insertion equilibrium ensembles. We observed an increase in HB occupancies between the NTP triphosphate tail and protein from initial binding to the inserted ensembles for ATP and dATP, with more protein-sugar HBs for the inserted ATP (with D623 and N691) than for the inserted dATP. On the other hand, the inserted RDV-TP ends up with fewer protein-triphosphate HBs than the inserted ATP but much stronger HBs (with T687 and N691) in the protein-sugar association. Meanwhile, GTP insertion led to many but weak (low occupancy) HBs formed, indicative of some instability.

Additionally (see **SI Figure S5**), in the open state, several protein residues (motif-F K551, K545, A558 and motif-B S682) form HBs with template +1 nt upon initial binding of GTP. While protein S501 (from motif G) forms HB with template +1 nt strongly in the presence of ATP (71%) and RDV-TP (78%), or marginally upon dATP (40%) and not present in GTP. In the closed state, the S501-template HB persists and becomes highly stabilized for every NTP insertion state. These findings again reflect distinctive HB patterns formed around the active site upon association of various NTPs, from initial binding to insertion. Nevertheless, due to various populations of NTP binding configurations, especially in the non-cognate initial binding state, it is not clear which HB interactions contribute to stabilize or destabilize cognate vs ncNTPs for nucleotide selectivity.

### 3.2 Constructing the PMFs for noncognate GTP/dATP from initial binding to insertion

Upon conformational samplings from the equilibrium ensemble simulations, we noticed essential variations of protein structural motifs along with diverse NTP dynamics. Accordingly, we included structural motifs and NTP configurations into constructing a collective reaction coordinate, based on RMSD changes with respective to the open and closed state [20, 21] structures (see **Methods**) [48,51]. Upon selecting appropriate reference states for each NTP initial binding/insertion system, we then proceeded to calculate the free energy profiles or PMFs of NTP insertion and to quantify the processes for NTP selectivity from initial binding (active site open) to the insertion (closed). The procedures of the PMF construction can be found in Methods. In current studies, we constructed the PMFs of the non-cognate GTP and dATP insertion, so that we can compare with the PMFs obtained previously for cognate ATP and RDV-TP [48].

From the constructed PMF, GTP upon initial binding displays stabilization in comparison with the insertion state, with Δ*G* = *G*_insertion_ *− G*_initial binding_ *∼*2 kcal/mol (*>* 0). This is in contrast with cognate ATP and RDV-TP analogue insertion, which are more stabilized in the insertion state, demonstrating Δ*G* (*<* 0) values of −5.2 kcal/mol and −2.7 kcal/mol, respectively (**Figure 3**) [48]. Nevertheless, the initially bound GTP forms no WB base pairing with the template nt+1, whereas approximately 81% WB base pairing between GTP and the template is identified in the insertion state. Hence, there have to be other interactions to stabilize the initial binding GTP (to be addressed in next subsection). Additionally, the PMF calculations show that the non-cognate GTP is subject to an insertion barrier *h^ins^ ∼*7 kcal/mol from the initial binding state, much larger than that of ATP and RDV-TP (*h^ins^ ∼*2.6 kcal/mol and 1.5 kcal/mol, respectively; see **Figure 3**) [48]. Note that the convergence of the PMF for GTP was reached after 50 ns per window in the umbrella sampling simulation, following the protocol without (or with) enforcing on the uracil template +1 nt (**SI Figure S6**), which show similar results.

**Figure 3:**
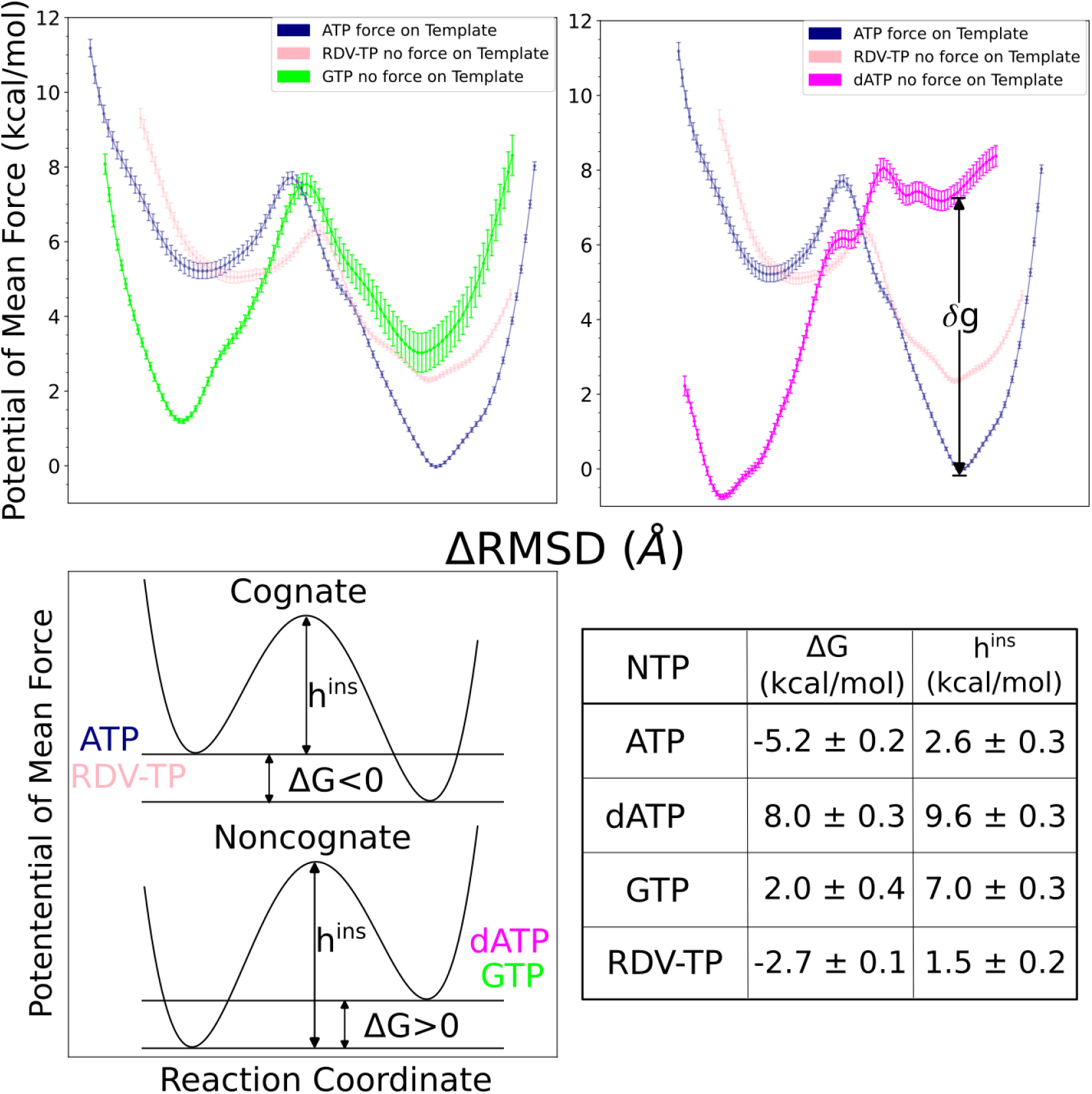
The potentials of mean force (PMFs) calculated for NTP from initial binding (active site open) to the insertion (closed) state via umbrella sampling simulations. The difference of RMSDs with respect to open and closed reference structures *RMSD*(*X, X*_Open Ref_) *− RMSD*(*X, X*_Closed Ref_) [48], was used as the reaction coordinate in the PMF construction (see Methods). Top left shows the PMF for GTP (green) and top right for that of dATP (magenta), both in comparison with the PMFs obtained for cognate ATP (blue) and analogue RDV-TP (pink) [48, 72]. Smoothing has been applied to each PMF for clarity (see original PMFs and converging tests in **Fig S6**). Note that the placement of the PMF of GTP relative to that of ATP/RDV-TP is according to the alchemical calculation in the closed state [72] while the placement of the PMF of dATP relative to that of ATP/RDV-TP is still uncertain, while the relative binding free energy at the closed state, denoted as *δg*, is estimated between 2-7 kcal/mol (see text). Bottom left shows schematics of the different PMF profiles for the cognate and ncNTP insertions. Bottom right shows the insertion free energy and barrier heights for the four PMFs shown on the top panel.

The placement of the GTP insertion PMF relative to that of ATP/RDV-TP was conducted according to alchemical simulations performed recently [72]. Given that the GTP-template WB base pairing geometries are comparatively stable in the insertion state (**Figure 3** *lower right*), we placed the GTP insertion state *∼*3 kcal/mol above that of the ATP insertion state, as the alchemical calculations indicate that ΔΔ*G*_binding_ 2.95 *±* 0.66 kcal/mol for the relative binding free energy of GTP with respect to ATP in the insertion state [72].

The dATP insertion PMF demonstrates significantly more stability in the initial binding state, with Δ*G ∼*8.0 kcal/mol comparing with the insertion state. Meanwhile, the insertion state of dATP can be hardly or only marginally stabilized. Besides, the non-cognate dATP is subject to an insertion barrier *h^ins^ ∼*9.6 kcal/mol, the highest among all NTP insertion cases examined (**Figure 3**). Similar to GTP, dATP does not form WC base pairing with the template nt+1 in the initial binding state but is capable of forming partial WC pairing with the template, though intermittently (64%), in the insertion state umbrella sampling windows. Therefore, there must also be some additional interactions stabilize or trapping dATP in the initial binding state, as reflected from the highly tilted PMF toward the initial binding state. Similar to GTP case, the constructed PMF of dATP reached convergence after 50 ns per window of the umbrella sampling simulation, following the protocol without (or with) force applied on the Uracil template (**Figure S6**), which both show similar results.

The placement of the dATP insertion PMF relative to that of ATP/RDV-TP is, however, less certain. Given a wide range of association configurations between dATP and template even in the insertion state (**Figure 2** *upper right*), one cannot directly use the alchemical calculation results, which were conducted around local configurational space (for stabilized ATP and slightly destabilized dATP) with limited sampling [72]. An estimation is nevertheless made here, based on relative binding free energy calculated locally between dATP and cognate ATP (*∼*2 kcal/mol) along with that from the mmPBSA calculation (*∼*7 kcal/mol) [72], suggesting a range of free energetic values 2-7 kcal/mol in between the inserted dATP and ATP (shown as parameter *δg* in **Figure 3**).

In order to further test the consistency of the PMF of insertion results for both GTP and dATP, we used SMD simulations to probe the stability of the insertion states for GTP (**Figure S7**) and dATP (**Figure S8**), respectively. To do that, the non-cognate GTP or dATP starting from the insertion state is pulled slightly away from the active site in the SMD simulations (see Methods), i.e., to start from the insertion state (active site closed) to the initial binding state (open) under the SMD force. In the case of GTP, it was robustly maintained within the insertion state without being able to cross the barrier (from closed to open) from three trials of the SMD simulations (one 500 ns, two 300 ns). In contrast, dATP was able to be readily pulled from the insertion state toward the initial binding state (closed to open) under the SMD force in all three simulations (300 ns each), demonstrating much lower stability or barrier from closed to open than that of GTP. These observations further support the instability of the dATP insertion state in comparison with the GTP insertion state, as being reflected from the PMFs constructed (**3**).

Next, we would proceed to examine additional interactions that stabilize or trap the non-cognate dATP or GTP in their initial binding state, i.e., as revealed from the PMFs constructed from the umbrella sampling simulation. However, before that, we want to examine whether the conformational space sampled between dATP/GTP and the template nt+1 near the initial binding (open) and insertion (closed) equilibrium in the umbrella sampling simulations overlap well with that from the equilibrium ensemble simulations. To do that, the NTP-template +1 nt base pairing geometries are compared between the equilibrium ensemble and the umbrella sampling simulations, from the latter three umbrella windows around the open/closed equilibrium were selected (40 ns RDV-TP/GTP/dATP and 160 ns ATP of simulation time per window). In **Figure 4**, one can see that the sampled geometries from the umbrella samplings are comparatively restricted, especially in the NTP initial binding state, but overlap well with the dominant configurations sampled from the equilibrium ensemble simulations, in particular, in the insertion state. For ATP and GTP, the initial binding configurations from the umbrella samplings overlap well with some stabilized population from the equilibrium ensemble, though deviations from the equilibrium ensemble show in the umbrella sampling case, indicating potentially the forcing impacts from the umbrella sampling simulations. For RDV-TP initial binding, the stable configuration of the RDV-TP-template in base stacking is well sampled, as the umbrella sampling path was launched from the base stacking configurations which leads to a low insertion barrier [48]. For the dATP initial binding, the umbrella sampling simulation also well covers one stabilized population. Hence, the significantly stabilized configuration of GTP/dATP upon initial binding, detected from the umbrella sampling simulations, appears to be well located within a subspace in the equilibrium ensemble.

**Figure 4:**
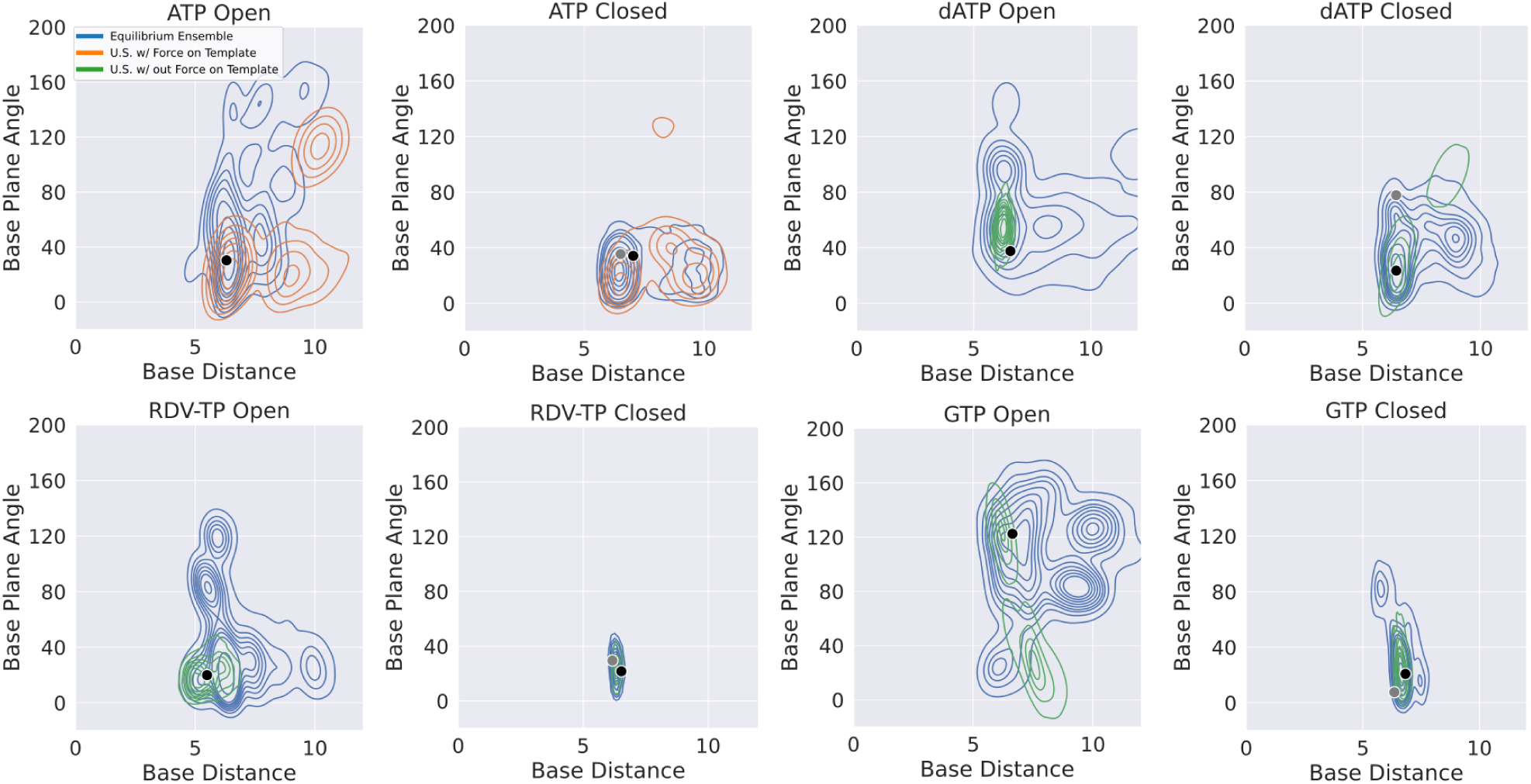
Comparing NTP-template association geometry distributions obtained from the umbrella sampling simulations (for the PMF calculations) and that from ensemble equilibrium simulations. A kernel density estimate has been used to visualize the data. Each simulation system (open and closed, for ATP, RDV-TP, dATP and GTP as in Figure 2), the equilibrium ensemble distribution is shown (blue), along with that obtained from the umbrella sampling (w/ force on template +1 nt in orange; w/out force on the template +1 nt in green; see **Methods** and **SI** for further details). The black dot indicates the reference state used to generate the initial paths (see **Methods**) for the umbrella sampling, and grey dot the reference state used in the alchemical calculations [72], for the full dataset see **Figure S9**.

Since the initially bound non-cognate dATP or GTP could not be well stabilized by association with the template +1 nt (nor 3’ end primer), it must be interactions from the RdRp protein along with the RNA scaffold around the active site that strongly hold the non-cognate dATP or GTP, which we examine and elaborate below.

### 3.3 Trapping noncognate dATP/GTP upon initial binding by persistent HB interactions from motif F/G/A, to NTP, template nt +1, or 3’ end primer

Though there were no WC or WB base pairing interactions observed for non-cognate dATP/GTP with the template uracil upon initial binding toward the open active site, some populations of dATP/GTP are strongly stabilized upon the initial binding according to their insertion free energetics or PMFs (shown in **Figure 3**). To gain understanding of this phenomena, we analyzed all HB interactions present among protein residues, NTP, and RNA strands (template and primer), around the active site. Given the variations amongst NTPs, we simplified the analyses by summing up overall HB populations exceeding 20% of occupancy (during the umbrella simulation 40ns/window) amongst the protein (residues within 10 Å of the active site center), template +1 nt, 3’ end of the primer, and the initially bound NTP; those HBs were further grouped according to interactions with the NTP on polyphosphate, sugar, or base (see **Figure 5**). The cumulative HB measure is then calibrated over that of the cognate ATP.

**Figure 5:**
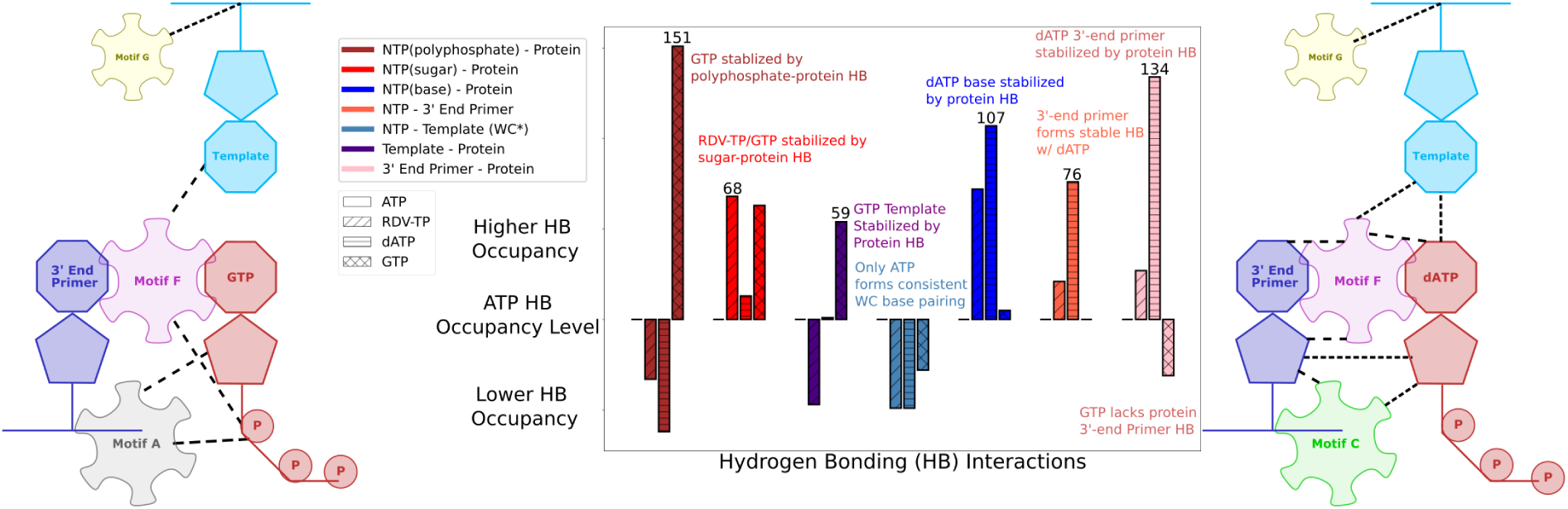
Summary of hydrogen bonding (HB) interactions that stabilize non-cognate GTP/dATP or surroundings upon initial binding from umbrella sampling simulations. Four interacting partners are considered: protein, incoming NTP (ATP, RDV-TP, dATP, and GTP), uracil template nucleotide +1, and the 3’-end primer. *Center:* The HBs with occupancy *>*20% in the simulations were identified among these four interaction partners. The relative HB occupancy levels are shown for the initial binding dATP and GTP (along with RDV-TP) with respect to that of cognate ATP as reference (with bar; see **SI Figure S10** for full HB statistics for all the simulation systems). Notable HB interactions that stabilize the non-cognate GTP and dATP initial binding systems are particularly denoted, respectively. Schematic and cartoon of key motifs stabilizing GTP (*left*) and dATP (*right*), along with template and 3’-end primer.

Notably, one finds that GTP initial binding is stabilized predominantly by HBs (and salt bridges in the case of positively charged residues LYS/ARG) formed between its polyphosphate group and the protein residues in motifs. In addition, the protein residues (motif F and G) stabilize particularly the template nt +1, due to the absence of WC or WB base pairing between the initially bound GTP and the template nt. Meanwhile, there is lack of protein HB association with the 3’ end primer around the initially bound GTP. However, this association is present around the initially bound ATP or RDV-TP, and the association appears even stronger around the non-cognate dATP initially bound.

In the case of dATP initial binding, though it does not form WC base pairing with the template nt+1, the adenine base is nonetheless stabilized via protein HB (again from motif-F). Intriguingly, despite dATP’s lack of a 2’ OH group, it still maintains a strong HB via the sugar of the 3’ end primer. As mentioned above, the 3’ end primer around the initially bound dATP displays the strongest HB interactions among all the NTP’s with the protein. Below, we show the individual HB interactions structurally around GTP and dATP and compare them with those in the case of cognate RDV-TP or ATP binding, respectively (**Figure 6** and **Figure 7**). One can find statistics of HBs formed between NTP (polyphosphate, sugar, or base) and protein residues or template nt +1/3’ end primer, and between protein residues and template nt+1 or the 3’ end primer in **SI Figure S10**. Notably, we have found that upon initial binding, GTP exhibits very strong HB or salt bridge interactions between motif-F K551/R553 (together with motif-A K621) and the polyphosphate (see **Figure 6A**). We attribute such interactions to hinder the insertion of GTP, or say, the phosphate-K551/R553 interactions contribute significantly to the barrier of GTP insertion. In the cognate initial binding, the ATP sugar forms a HB with motif-C D760. In contrast, GTP initial binding is mainly stabilized by HB between sugar and motif-A D623 (from umbrella sampling simulations). Instead of WB pairing with the non-cognate GTP upon initial binding, the template nt+1 base forms HBs with motif-F K545 and A558 (see **Figure 6B**). Furthermore, the template nt +1 backbone forms HB with motif-G K511, as opposed to motif-G S501 seen in the other NTP binding cases. Overall, the protein motifs F and G seem to well stabilize the template nt +1 upon the base mismatched GTP binding.

**Figure 6:**
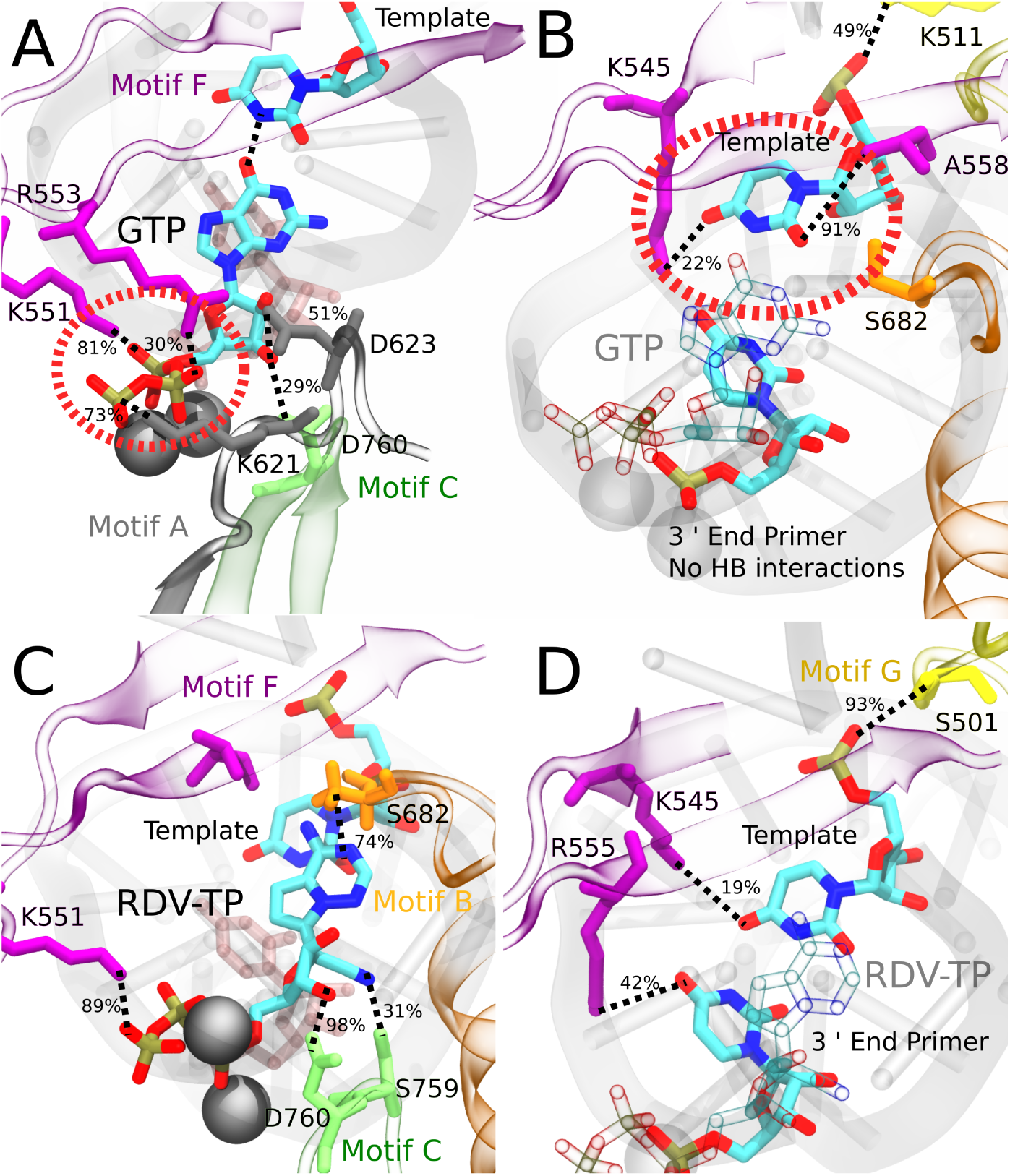
Comparing key interactions that stabilize the non-cognate GTP (and surroundings) and drug analogue RDV-TP in the initial binding system. The conserved protein motifs are shown in cartoon representation while interacting residues from these motifs are shown in licorice using the same color. A/C: Incoming GTP/RDV-TP and template nucleotide are shown in licorice colored by atom name. Hydrogen bonds (HBs) formed between GTP (or RDV-TP) and template nucleotide uracil/protein/3’ end RNA primer. Orange circle highlights the essential interactions involved in ‘trapping’ non-cognate GTP phosphate in the initially bound state (A), which are absent for RDV-TP phosphate (C). B/D: Incoming GTP/RDV-TP is shown in transparent representation for clarity, 3’-end primer and template nucleotide are shown in licorice colored by atom name. HBs formed between the protein and template for stabilization, without proper WB base pairing for GTP-template (B) and with template base stacking in RDV-TP (D). For detailed HB occupancy plots see **SI Figure S10**

**Figure 7:**
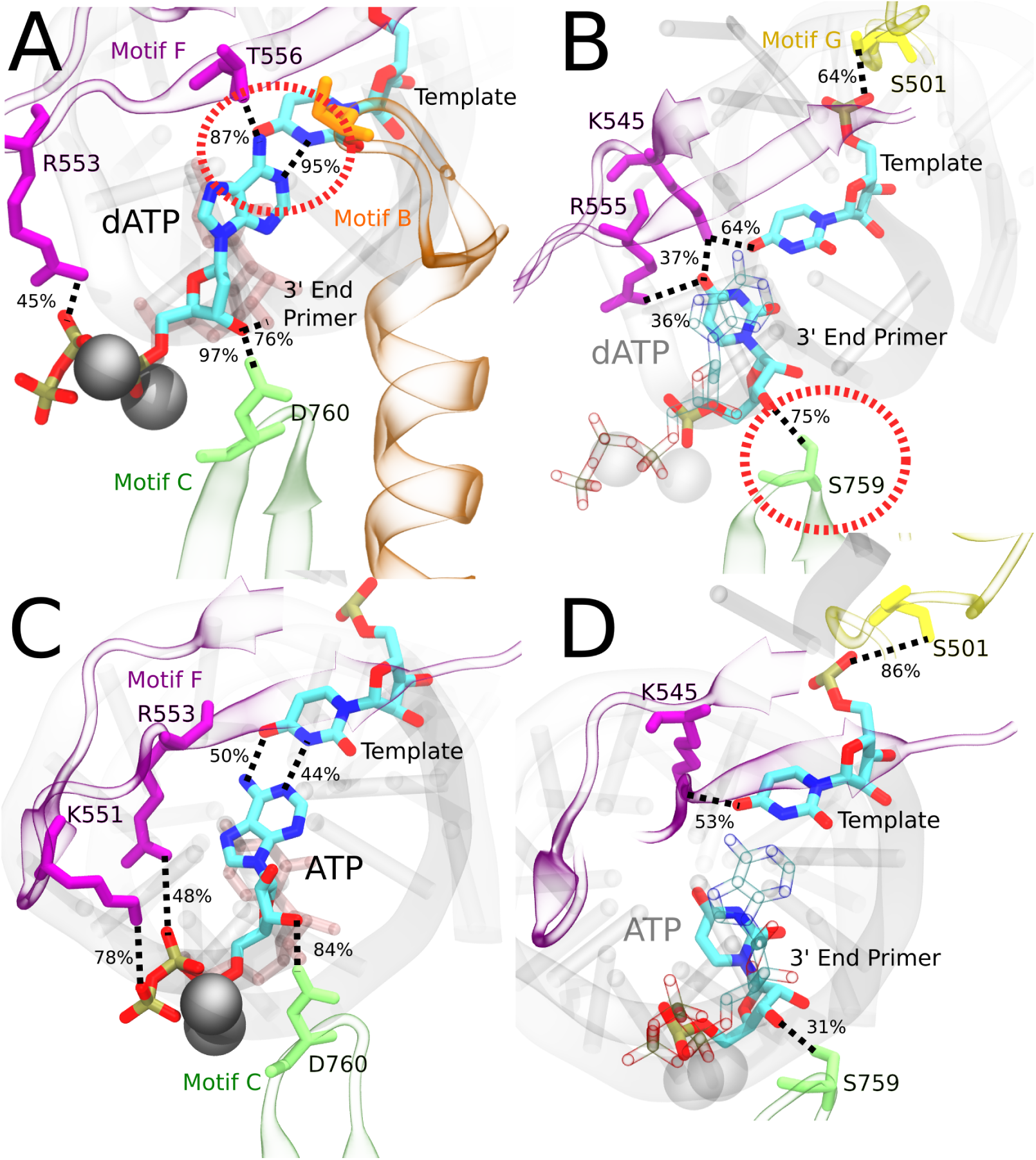
Comparing the key interactions that stabilize the non-cognate dATP (and surroundings) and cognate ATP in the initial binding system. The conserved protein motifs are shown in cartoon representation while interacting residues from these motifs are shown in licorice using the same color. A/C: Incoming dATP/ATP and template nucleotide are shown in licorice colored by atom name. Hydrogen bonds (HBs) formed between dATP (or ATP) and template nucleotide uracil/protein/3’-end RNA primer. Orange circle highlights the strongest interactions involved in ‘trapping’ non-cognate dATP in the initially bound state. B/D: Incoming dATP/ATP is shown in transparent representation for clarity, 3’-end primer and template nucleotide are shown in licorice colored by atom name. HBs formed between the protein and template nucleotide uracil / 3’-end RNA primer for stabilization, in the absence of proper WC base pairing in dATP (B). HBs formed between the protein and template nucleotide uracil / 3’-end RNA primer for stabilization, with proper WC base pairing in ATP (D). For detailed HB occupancy plots see **SI Figure S10**.

In the case of RDV-TP initial binding via base stacking with the template nt +1 (in the absence of force on the template nt), however, only one HB is observed on the polyphosphate from motif-F K551 (see **Figure 6C**), so that the RDV-TP won’t be hindered by the phosphate interaction for its insertion [48]. In addition, the base stacking configuration of RDV-TP with the template allows for a unique HB to form between motif-B S682 and the RDV-TP base, while the template nt +1 has fewer HBs with motif-F than that upon GTP initial binding (**Figure 6D**).

In comparison, as from current umbrella sampling simulations, dATP upon initial binding forms a single HB with template nt +1 at a very high occupancy of 95%. In addition, a HB is uniquely established between the dATP base and motif-F T556 at an occupancy of 87% (**Figure 7A**). The dATP initial binding also forms two persistent HBs between the sugar and motif-C D760 and the 3’ end primer, respectively. The template nt +1 is further stabilized by interactions with motif-F K545 and S501G, as seen with the cognate initial binding (**Figure 7B**). Importantly, 3’ end primer forms the most persistent HBs with motif-F K545/R555 and motif-C S759 in the case of dATP initial binding.

In the case of ATP initial binding, its base is stabilized by forming variable WC base pairing. Note that the initially bound ATP formed two HBs with template nt +1, with occupancies of 50% and 44%, respectively. The ATP sugar forms a consistent single HB with motif-C D760. Stable associations also form between motif-F K551/R553/R555 and the ATP polyphosphate (**Figure 7C**). Such interactions were suggested to facilitate the cognate ATP insertion instead [48]. The template nt +1 forms stable HB with motif-F K545 and motif-G S501, similar to dATP initial binding. In addition, the 3’ end primer forms only a single transient HB with motif-C S759 (**Figure 7D**), weaker than that is present between the 3’ end and motif-C/F upon the dATP initial binding (**Figure 7B**).

## 4 DISCUSSION

In current work, we have focused on computationally probing from initial binding to the insertion and selectivity mechanisms of noncognate natural nucleotides to SARS-CoV-2 RdRp. The insertion step process involves subtle but still substantial conformational change of the RdRp pol domain (**Figure 1**), leading to essentially an active site open or nucleotide initial binding state to the active site closed or insertion state [31], with coordination of seven highly conserved structural motifs. In all NTP incorporation systems simulated, the fingers subdomain displays similar conformational flexibility as the palm subdomain, which moves closer to the finger’s subdomain in the insertion state than in the initial NTP binding state (**Figure S1**). In the insertion state of cognate ATP or RDV-TP, motif-B and C have similar conformational flexibility (via RMSD) demonstrated in the equilibrium simulation, while this feature is absent in noncognate GTP and dATP (Figure S2). Overall, the motifs respond differently for each incoming NTP studied. The equilibrium ensemble simulations showed generically that the insertion state sampled a restricted subspace between the NTP and template nt +1 over that of the initial binding state, which indeed accommodates a wide range of configurations (**Figure 2**). Additionally, RNA template/primer nucleotides or protein residues around the active site forms more HBs with the NTP in the insertion state than in the initial binding state (**Figure S4**). Due to time scale limit of equilibrium sampling, we probed the NTP insertion dynamics and calculated the corresponding insertion energetics using the umbrella sampling methods. The energetic profile or insertion PMF was constructed along a collective RC according to a difference of RMSDs between the modeled intermediate structure of the RdRp and the open and closed reference states, respectively. The essential atomic coordinates included those of backbone atoms from seven highly conserved structural motifs (A to G), which are crucial for recruiting nucleotide substrates with selectivity and supporting catalysis [73], and heavy atoms on incoming NTP along with (or without) the template nt +1. While such a choice on enforcing on the template nt or not played some significant role in the PMF construction of cognate ATP and analogue RDV-TP (see **Figure S10**) [48], it made little difference in the non-cognate dATP or GTP results, e.g., as observed from the base pairing geometry measured between NTP and template nt +1 (**Figure 4&S9**). The insertion barriers were not affected much by the choices for dATP or GTP either (Figure S6). In contrast with the insertion PMFs of cognate ATP and RDV-TP analogue that bias toward a more stabilized insertion state, we have found intriguingly that the initial binding states for ncNTPs (dATP and GTP currently) can be much more stabilized than their insertion state. In addition, the insertion barrier of ncNTP becomes very high (up to 7-10 kcal/mol), also in contrast with the marginally low insertion barriers of cognate ATP and RDV-TP (*∼*2 kcal/mol) identified previously [48]. Such free energetic calculations and structural dynamics examinations reveal intrinsic nucleotide selectivity of SARS-CoV-2 RdRp, i.e., to inhibit the insertion of ncNTPs to the active site by trapping the ncNTPs *off-path* upon initial binding to the peripheral of the RdRp active site.

### 4.1 Free energetics favor insertion of cognate NTPs but disfavor insertion of non-cognate NTPs

In previous work [48], we calculated the insertion PMFs for ATP and RDV-TP, respectively. While the RDV-TP initial binding could form WC base pairing with the template nt+1, a more stabilized conformation was found for RDV-TP in base stacking with the template nt +1. Besides, the RDV-TP insertion barrier would become high (*h^ins^ ∼*5 kcal/mol) when the enforcing in the umbrella sampling simulation included the template nt +1 (**Figure S11A**). The striking feature was due to the enhanced HB (or salt-bridge from positively charged LYS/ARG) interactions between the motif-F residues (K551/R553/R555) and the RDV-TP polyphosphate (**Figure S11B**), upon the enforcing on the template nt. Removing forcing on the template nt+1, i.e., allowing sufficient thermal fluctuations on the template, however, the motif-F interactions with the polyphosphate reduce, and the insertion barrier lowers to a marginal value (*h^ins^ ∼*1.5 kcal/mol).

Upon the cognate ATP binding, including the template nt +1 for enforcing in the umbrella sampling simulations nevertheless supports an insertion barrier *h^ins^* as low as *∼*2.6 kcal/mol (**Figure S11A**). The well controlled template nt+1 facilitated WC base pairing and supported enhanced HB interactions between the motif-F K551/R555 and polyphosphate (**Figure S11C**). Consequently, it was suggested that the motif-F interactions with the phosphate facilitate insertion of the cognate ATP, while the motif F-phosphate interactions also appear to be important for NTP insertion in the PV RdRp [74]. Regardless of the exact protocol, the insertion state was always more stable than the initial binding state for the cognate ATP and analogue RDV-TP, both of which are actively biased or recruited into the closed active site for catalytic incorporation to the 3’-end of primer, experiencing marginally high barriers of insertion to the active site [48].

In contrast, upon the initial binding of ncNTP (dATP or GTP), a large configuration space of the ncNTP with respect to the template+1 nt and 3’-end primer was identified, and the PMF always tilts toward the initial binding state, i.e., biased or energetically stabilized upon initial binding of certain ncNTP configurations (**Figure 3&S6**). Additionally, the insertion barriers insertion become very large for noncognate GTP and dATP (*h^ins^ ∼*7.0 and 9.6 kcal/mol, respectively). The stability bias toward the initial binding state and tremendously large barrier of insertion seem to trap the ncNTP upon initial binding at certain configurations (or the *off-path*), which would likely lead to dissociation of the ncNTPs from the active site or from the RdRp in the end.

Our current discoveries on such stabilization of the non-cognate substrate upon initial binding may seem counterintuitive. Commonly, high binding affinity of a ligand substrate to the receptor protein indicates a preference of the receptor to the substrate [75, 76]. Such an idea prevails in the drug design which aims at identifying high-affinity binders. In the current viral RdRp system, however, the NAC proceeds with two pre-chemical steps: substrate initial binding and insertion. As shown in current study, a high substrate binding affinity at the first step may also contribute to high insertion barrier for the second step, which slows down the NAC or an enzymatic cycle. Hence, the ncNTP stabilization or trapping upon initial binding becomes an intriguing but effective strategy to deter or inhibit the non-cognate substrate from further incorporation.

### 4.2 Key residues from conserved motifs detect and stabilize the ncNTP and its surroundings at the initial binding off-path state

Current free energy calculations reveal that the noncognate GTP/dATP is more stabilized upon initial binding to the RdRp active site than in the insertion state, in contrast with cognate ATP/RDV-TP to be more stabilized in the insertion state. To explain notable stabilized configurations sampled for the ncNTP initial binding state from the umbrella sampling simulations, we identified a variety of HB interactions around the active binding site formed among NTP (base, sugar and phosphate), RNA strands (template and primer), and the conserved protein motifs (**Figure 5**). To well explain the trapping mechanism, we can also compare current system with a previously studied RdRp from Enterovirus or EV [31], which is structurally similar to SARS CoV-2 RdRp. In EV RdRp, NTP insertion to the active site is suggested via several steps: first the base recognition, next the ribose sugar recognition, and then followed by the active site open to closed conformational transition, accompanied by the palm subdomain (motifs A,B,C,D,E) closing. Below we connect current observations of NTP initial binding to those suggested steps.

In the case of GTP initial binding, the template nt +1 (uracil) fails to interact with the mismatch GTP in the absence of WC or WB base pairing. The template nt +1 and GTP cannot be mutually stabilized, hence the protein motif-F residues (K545/A558) respond by stabilizing the template base. Additionally, motif-G K511 forms HB with the template RNA backbone, instead of S501 in the ATP/RDV-TP system. Given the non-stabilized GTP base, the sugar is next checked by the protein via motif-A D623, which is prominent upon GTP initial binding (in the umbrella sampling ensemble). The D623 interaction brings motif-A closer to GTP and allows a unique HB (or salt bridge) from K621 to the polyphosphate. Substantial HBs (or salt bridges) with the polyphosphate come additionally from motif-F K551/R553, similarly as seen in RDV-TP (with force on template nt +1). In both cases of initial binding (GTP and RDV-TP with forcing on the template nt+1), the insertion barriers are high. Such observations support the previously proposed mechanism that the Lys/Arg interactions with phosphates inhibit the ncNTP insertion but facilitate cognate NTP insertion [48], or say, the protein-NTP phosphate interactions play a significant role in nucleotide selectivity or fidelity control. Recent NMR experiments on the structurally similar PV RdRp suggest that interactions from charged residues in motif-F are an important fidelity checkpoint, as they allow the triphosphate to rearrange prior to catalysis [77].

In the case of dATP binding, since dATP has the same base as the cognate ATP, it should be capable of forming stable WC base pairing with the template nt, but it does not. Additionally, dATP fails on the sugar recognition. The sugar is unable to anchor in the active site due to missing the 2’ OH functional group. Instead, the 3’ OH group forms HB with motif-C D760, similar to ATP and RDV-TP. Meanwhile, a unique HB forms between the dATP sugar and 3’ end primer HB. As a result, dATP base adopts a tilted conformation, which supports only a single persistent HB w/ template nt +1 (**Figure 7A**). The dATP base is then further stabilized by motif-F T556. The missing WC base pairing between dATP and the template nt +1 in is supplemented by another HB formed between the dATP base and motif-F K545. Additional stabilization of dATP initial binding comes from the HB formed between the 3’ end primer from motif-C S759, and the 3’-end primer further forms HB with dATP sugar. The dATP sugar further forms HB with D760 from motif C. Hence, it appears that the 3’-end primer plays an important role in stabilizing dATP with a network of HB interactions upon initial binding or say, *off-path*.

Although the non-cognate dATP initial binding stability appears perplexing, prior crystal structure studies on the structurally similar PV RdRp have used dCTP to stall and resolve the RdRp structure in the open state [30], supporting a stable binding configuration of dNTP binding to the viral RdRp. In addition, experimental work on the nsp14 exonuclease enzyme of SARS-CoV-2 has shown that for excision of an incorrect nt, the nt needs to have a proper RNA sugar, 2’ and 3’ OH functional group [78], indicating that the enzyme may require some nucleotide selectivity to prevent dNTP’s from chemical incorporation. Furthermore, recent experimental studies have shown that an elongation complex soaked in solution with dNTPs had no catalytic activity [79]. Michaelis–Menten kinetics also showed the SARS-CoV-2 RdRp selectivity of ATP over dATP is *∼*1000 (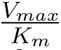 = 0.02 in dATP and 23 in ATP) [80]. Similar trends were observed in PV *in vitro* biochemical studies with *∼*117 discrimination factor 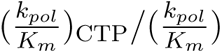dCTP [81]. Other computational works also tested the design of inhibitors with ribose sugar modifications or removal of the OH function groups entirely [36, 37].

## 5 Conclusion

To conclude, we have employed all-atom MD simulations and demonstrated intrinsic or natural nucleotide selectivity of SARS-CoV-2 RdRp, in which the ncNTP is well stabilized or trapped upon initial binding to certain off-path configurations, as the highly conserved structural motifs F/G/A/C of the viral RdRp form HBs with ncNTP, RNA template nt, and/or 3’-end RNA primer. Intrinsically, it is not the polymerase enzyme that determines a right/cognate or wrong/non-cognate nucleotide substrate in the template-based polymerization. The cognate or non-cognate NTP species are determined by the RNA template nt primarily via the WC base pairing, and additionally by the 3’-end primer via base stacking etc. With incoming ncNTP that is incapable of stabilizing the template counterpart or the 3’-end primer, the RdRp structural motifs sense such instability, and then takes over to select against the ncNTP. Presumably, ncNTP can be rejected in case of low binding affinity, i.e., upon initial binding to RdRp. However, it appears that in SARS-CoV-2 RdRp, the ncNTP can be particularly stabilized, say *off-path*, upon initial binding to certain configurations, so that to be prevented or inhibited from further insertion to the active site. Such mechanism of nucleotide selection seems to be well supported by the two-step binding and insertion processes, *pre-chemically*, in the single-subunit viral polymerase enzymes. Partial off-path initial binding and inhibition for insertion of ncNTPs were also suggested computationally for single-subunit T7 RNAP [51, 82].

Technically, we have employed the umbrella sampling simulations to characterize the slow NTP insertion dynamics that is accompanied by the open to closed conformational changes around the RdRp active site. As the slow pre-chemical conformational transition likely takes place over milliseconds, it becomes indispensable to enhance computational sampling while limiting artifacts to be introduced. In the simulation, we well manipulated collective atomic coordinates from all structural motifs along with key players of NTP/template. Nevertheless, how to identify the most essential coordinates is a continuous challenging issue [83, 84].

Additionally, accelerated simulations [85] may be conducted to sample a sufficient amount of conformational transitions, i.e., from open to closed. For multiple NTP species incorporation, enhanced computational sampling would become even more demanding, considering that multiple reaction paths exist [86]. Further exploration and validation of our current studies would require substantial experimental studies. Resolving high-resolution structures of the SARS-CoV-2 RdRp complexes with the stabilized *off-path* initial binding configurations of non-cognate dATP or GTP, as being proposed in current computational work, would be highly anticipated.

## Supporting information

Sample Input Files

## ACKNOWLEDGEMENTS

Part of this work was supported by National Science Foundation (NSF) Award #2028935. This research used resources of the Oak Ridge Leadership Computing Facility, which is a DOE Office of Science User Facility supported under Contract DE-AC05-00OR22725. Additional computational resources were provided by UCI HPC3 - High Performance Community Computing Cluster. JY has also been supported by the CMCF of UCI via NSF DMS 1763272 and the Simons Foundation grant #594598 and start-up funding from UCI.

## 6 Data Availability

Input files available with provided zip. Trajectories available on request.

## References

[1] Harvey, W. T., Carabelli, A. M., Jackson, B., Gupta, R. K., Thomson, E. C., Harrison, E. M., Ludden, C., Reeve, R., Rambaut, A., Peacock, S. J., and Robertson, D. L. (2021) SARS-CoV-2 variants, spike mutations and immune escape. Nature Reviews Microbiology 2021 19:7, 19(7), 409–424.

[2] Araf, Y., Akter, F., Tang, Y. d., Fatemi, R., Parvez, M. S. A., Zheng, C., and Hos-sain, M. G. (2022) Omicron variant of SARS-CoV-2: Genomics, transmissibility, and responses to current COVID-19 vaccines. Journal of Medical Virology, 94(5), 1825–1832.

[3] Pandey, P., Rane, J. S., Chatterjee, A., Kumar, A., Khan, R., Prakash, A., and Ray, S. (2020) Targeting SARS-CoV-2 spike protein of COVID-19 with naturally occurring phytochemicals: an in silico study for drug development. Journal of Biomolecular Structure and Dynamics, 39(16), 1–11.

[4] Papageorgiou, A. C. and Mohsin, I. (2020) The SARS-CoV-2 Spike Glycoprotein as a Drug and Vaccine Target: Structural Insights into Its Complexes with ACE2 and Antibodies. Cells 2020, Vol. 9, Page 2343, 9(11), 2343.

[5] Almehdi, A. M., Khoder, G., Alchakee, A. S., Alsayyid, A. T., Sarg, N. H., and Soliman, S. S. (2021) SARS-CoV-2 spike protein: pathogenesis, vaccines, and potential therapies. Infection, 49(5), 855–876.

[6] Ullrich, S. and Nitsche, C. (2020) The SARS-CoV-2 main protease as drug target. Bioorganic and Medicinal Chemistry Letters, 30(17), 127377.

[7] Jin, Z., Wang, H., Duan, Y., and Yang, H. (2021) The main protease and RNA-dependent RNA polymerase are two prime targets for SARS-CoV-2. Biochemical and Biophysical Research Communications, 538, 63–71.

[8] Tarannum, H., Rashmi, K., and Nandi, S. (2021) Exploring the SARS-Cov-2 Main Protease (Mpro) and RdRp Targets by Updating Current Structure-based Drug Design Utilizing Co-crystals to Combat COVID-19. Current Drug Targets, 23(8), 802–817.

[9] Magazine, N., Zhang, T., Wu, Y., McGee, M. C., Veggiani, G., and Huang, W. (2022) Mutations and Evolution of the SARS-CoV-2 Spike Protein. Viruses 2022, Vol. 14, Page 640, 14(3), 640.

[10] Papanikolaou, V., Chrysovergis, A., Ragos, V., Tsiambas, E., Katsinis, S., Manoli, A., Papouliakos, S., Roukas, D., Mastronikolis, S., Peschos, D., Batistatou, A., Kyrodimos, E., and Mastronikolis, N. (2022) From delta to Omicron: S1-RBD/S2 mutation/deletion equilibrium in SARS-CoV-2 defined variants. Gene, 814, 146134.

[11] Mótyán, J. A., Mahdi, M., Hoffka, G., and Tőzsér, J. (2022) Potential Resistance of SARS-CoV-2 Main Protease (Mpro) against Protease Inhibitors: Lessons Learned from HIV-1 Protease. International Journal of Molecular Sciences 2022, Vol. 23, Page 3507, 23(7), 3507.

[12] Iketani, S., Mohri, H., Culbertson, B., Hong, S. J., Duan, Y., Luck, M. I., Annavajhala, M. K., Guo, Y., Sheng, Z., Uhlemann, A. C., Goff, S. P., Sabo, Y., Yang, H., Chavez, A., and Ho, D. D. (2022) Multiple pathways for SARS-CoV-2 resistance to nirmatrelvir. Nature 2022 613:7944, 613(7944), 558–564.

[13] Hu, Y., Lewandowski, E. M., Tan, H., Zhang, X., Morgan, R. T., Zhang, X., Jacobs, L. M. C., Butler, S. G., Gongora, M. V., Choy, J., Deng, X., Chen, Y., and Wang, J. (2022) Naturally occurring mutations of SARS-CoV-2 main protease confer drug resistance to nirmatrelvir. bioRxiv, p. 2022.06.28.497978.

[14] Ahn, D. G., Choi, J. K., Taylor, D. R., and Oh, J. W. (2012) Biochemical characterization of a recombinant SARS coronavirus nsp12 RNA-dependent RNA polymerase capable of copying viral RNA templates. Archives of Virology, 157(11), 2095–2104.

[15] Sevajol, M., Subissi, L., Decroly, E., Canard, B., and Imbert, I. (2014) Insights into RNA synthesis, capping, and proofreading mechanisms of SARS-coronavirus. Virus Research, 194, 90–99.

[16] Zhu, W., Chen, C. Z., Gorshkov, K., Xu, M., Lo, D. C., and Zheng, W. (2020) RNA-Dependent RNA Polymerase as a Target for COVID-19 Drug Discovery. SLAS Discovery, 25(10), 1141–1151.

[17] Grellet, E., L’Hô te, I., Goulet, A., and Imbert, I. (2022) Replication of the coronavirus genome: A paradox among positive-strand RNA viruses. Journal of Biological Chemistry, 298(5), 1–15.

[18] Gong, P. (2022) Within and Beyond the Nucleotide Addition Cycle of Viral RNA-dependent RNA Polymerases. Frontiers in Molecular Biosciences, 8, 822218.

[19] Long, C., Romero, M. E., La Rocco, D., and Yu, J. (2021) Dissecting nucleotide selectivity in viral RNA polymerases. Computational and Structural Biotechnology Journal, 19, 3339–3348.

[20] Gao, Y., Yan, L., Huang, Y., Liu, F., Zhao, Y., Cao, L., Wang, T., Sun, Q., Ming, Z., Zhang, L., Ge, J., Zheng, L., Zhang, Y., Wang, H., Zhu, Y., Zhu, C., Hu, T., Hua, T., Zhang, B., Yang, X., Li, J., Yang, H., Liu, Z., Xu, W., Guddat, L. W., Wang, Q., Lou, Z., and Rao, Z. (2020) Structure of the RNA-dependent RNA polymerase from COVID-19 virus. Science, 368(6492), 779–782.

[21] Yin, W., Mao, C., Luan, X., Shen, D.-D., Shen, Q., Su, H., Wang, X., Zhou, F., Zhao, W., Gao, M., Chang, S., Xie, Y.-C., Tian, G., Jiang, H.-W., Tao, S.-C., Shen, J., Jiang, Y., Jiang, H., Xu, Y., Zhang, S., Zhang, Y., and Xu, H. E. (2020) Structural basis for inhibition of the RNA-dependent RNA polymerase from SARS-CoV-2 by remdesivir. Science, 368(6498), 1499–1504.

[22] Hillen, H. S., Kokic, G., Farnung, L., Dienemann, C., Tegunov, D., and Cramer, P. (2020) Structure of replicating SARS-CoV-2 polymerase. Nature, 584(7819), 154–156.

[23] Chen, J., Malone, B., Llewellyn, E., Grasso, M., Shelton, P. M., Olinares, P. D. B., Maruthi, K., Eng, E. T., Vatandaslar, H., Chait, B. T., Kapoor, T. M., Darst, S. A., and Campbell, E. A. (2020) Structural Basis for Helicase-Polymerase Coupling in the SARS-CoV-2 Replication-Transcription Complex. Cell, 182(6), 1560–1573.

[24] Wang, Q., Wu, J., Wang, H., Gao, Y., Liu, Q., Mu, A., Ji, W., Yan, L., Zhu, Y., Zhu, C., Fang, X., Yang, X., Huang, Y., Gao, H., Liu, F., Ge, J., Sun, Q., Yang, X., Xu, W., Liu, Z., Yang, H., Lou, Z., Jiang, B., Guddat, L. W., Gong, P., and Rao, Z. (2020) Structural Basis for RNA Replication by the SARS-CoV-2 Polymerase. Cell, 182(2), 417–428.

[25] Malone, B., Chen, J., Wang, Q., Llewellyn, E., Choi, Y. J., Olinares, P. D. B., Cao, X., Hernandez, C., Eng, E. T., Chait, B. T., Shaw, D. E., Landick, R., Darst, S. A., and Campbell, E. A. (2021) Structural basis for backtracking by the SARS-CoV-2 replication-transcription complex. Proceedings of the National Academy of Sciences of the United States of America, 118(19), 2102516118.

[26] Bravo, J. P., Dangerfield, T. L., Taylor, D. W., and Johnson, K. A. (2021) Remdesivir is a delayed translocation inhibitor of SARS-CoV-2 replication. Molecular Cell, 81(7), 1548–1552.

[27] Peng, Q., Peng, R., Yuan, B., Wang, M., Zhao, J., Fu, L., Qi, J., and Shi, Y. (2021) Structural Basis of SARS-CoV-2 Polymerase Inhibition by Favipiravir. Innovation, 2(1).

[28] Naydenova, K., Muir, K. W., Wu, L. F., Zhang, Z., Coscia, F., Peet, M. J., Castro-Hartmann, P., Qian, P., Sader, K., Dent, K., Kimanius, D., Sutherland, J. D., Löowe, J., Barford, D., and Russo, C. J. (2021) Structure of the SARS-CoV-2 RNA-dependent RNA polymerase in the presence of favipiravir-RTP. Proceedings of the National Academy of Sciences of the United States of America, 118(7), e2021946118.

[29] Malone, B. F., Perry, J. K., Olinares, P. D. B., Lee, H. W., Chen, J., Appleby, T. C., Feng, J. Y., Bilello, J. P., Ng, H., Sotiris, J., Ebrahim, M., Chua, E. Y., Mendez, J. H., Eng, E. T., Landick, R., Göotte, M., Chait, B. T., Campbell, E. A., and Darst, S. A. (2023) Structural basis for substrate selection by the SARS-CoV-2 replicase. Nature, 614(7949), 781–787.

[30] Gong, P. and Peersen, O. B. (2010) Structural basis for active site closure by the poliovirus RNA-dependent RNA polymerase. Proceedings of the National Academy of Sciences of the United States of America, 107(52), 22505–22510.

[31] Shu, B. and Gong, P. (2016) Structural basis of viral RNA-dependent RNA polymerase catalysis and translocation. Proceedings of the National Academy of Sciences, 113(28), E4005–E4014.

[32] Beigel, J. H., Tomashek, K. M., Dodd, L. E., Mehta, A. K., Zingman, B. S., Kalil, A. C., Hohmann, E., Chu, H. Y., Luetkemeyer, A., Kline, S., Lopez de Castilla, D., Finberg, R. W., Dierberg, K., Tapson, V., Hsieh, L., Patterson, T. F., Paredes, R., Sweeney, D. A., Short, W. R., Touloumi, G., Lye, D. C., Ohmagari, N., Oh, M.-d., Ruiz-Palacios, G. M., Benfield, T., Fätkenheuer, G., Kortepeter, M. G., Atmar, R. L., Creech, C. B., Lundgren, J., Babiker, A. G., Pett, S., Neaton, J. D., Burgess, T. H., Bonnett, T., Green, M., Makowski, M., Osinusi, A., Nayak, S., and Lane, H. C. (2020) Remdesivir for the Treatment of Covid-19 — Final Report. New England Journal of Medicine, 383(19), 1813–1826.

[33] Warren, T. K., Jordan, R., Lo, M. K., Ray, A. S., Mackman, R. L., Soloveva, V., Siegel, D., Perron, M., Bannister, R., Hui, H. C., Larson, N., Strickley, R., Wells, J., Stuthman, K. S., Van Tongeren, S. A., Garza, N. L., Donnelly, G., Shurtleff, A. C., Retterer, C. J., Gharaibeh, D., Zamani, R., Kenny, T., Eaton, B. P., Grimes, E., Welch, L. S., Gomba, L., Wilhelmsen, C. L., Nichols, D. K., Nuss, J. E., Nagle, E. R., Kugelman, J. R., Palacios, G., Doerffler, E., Neville, S., Carra, E., Clarke, M. O., Zhang, L., Lew, W., Ross, B., Wang, Q., Chun, K., Wolfe, L., Babusis, D., Park, Y., Stray, K. M., Trancheva, I., Feng, J. Y., Barauskas, O., Xu, Y., Wong, P., Braun, M. R., Flint, M., McMullan, L. K., Chen, S. S., Fearns, R., Swaminathan, S., Mayers, D. L., Spiropoulou, C. F., Lee, W. A., Nichol, S. T., Cihlar, T., and Bavari, S. (2016) Therapeutic efficacy of the small molecule GS-5734 against Ebola virus in rhesus monkeys. Nature, 531(7594), 381–385.

[34] Ahmed, S., Mahtarin, R., Islam, S., Das, A., Al Mamun, S., Samina, A., Ali, A., Das, S., Al Mamun, A., and Ahmed, S. (2022) Remdesivir analogs against SARS-CoV-2 RNA-dependent RNA polymerase. Journal of Biomolecular Structure and Dynamics, 40(21), 11111–11124.

[35] Koulgi, S., Jani, V., Mallikarjunachari Uppuladinne, V. N., Sonavane, U., and Joshi, R. (2022) Structural insight into the binding interactions of NTPs and nucleotide analogues to RNA dependent RNA polymerase of SARS-CoV-2. Journal of Biomolecular Structure and Dynamics, 40(16), 7230–7244.

[36] Parise, A., Ciardullo, G., Prejanò, M., De La Lande, A., and Marino, T. (2022) On the Recognition of Natural Substrate CTP and Endogenous Inhibitor ddhCTP of SARS-CoV-2 RNA-Dependent RNA Polymerase: A Molecular Dynamics Study. Journal of Chemical Information and Modeling, 62(20), 4916–4927.

[37] Li, Y., Zhang, D., Gao, X., Wang, X., and Zhang, L. (2022) 2- and 3-Ribose Modifications of Nucleotide Analogues Establish the Structural Basis to Inhibit the Viral Replication of SARS-CoV-2. Journal of Physical Chemistry Letters, 13, 4111–4118.

[38] Giannetti, M., Mazzuca, C., Ripani, G., and Palleschi, A. (2023) Inspection on the Mechanism of SARS-CoV-2 Inhibition by Penciclovir: A Molecular Dynamic Study. Molecules, 28(1), 191.

[39] Bera, S. C., Seifert, M., Kirchdoerfer, R. N., van Nies, P., Wubulikasimu, Y., Quack, S., Papini, F. S., Arnold, J. J., Canard, B., Cameron, C. E., Depken, M., and Dulin, D. (2021) The nucleotide addition cycle of the SARS-CoV-2 polymerase. Cell Reports, 36(9).

[40] Luo, X., Xu, T., Gao, X., and Zhang, L. (2022) Alternative role of motif B in template dependent polymerase inhibition. Chinese Journal of Chemical Physics, 35(3), 407–412.

[41] Yuzenkova, Y., Bochkareva, A., Tadigotla, V. R., Roghanian, M., Zorov, S., Severinov, K., and Zenkin, N. (2010) Stepwise mechanism for transcription fidelity. BMC Biology, 8(1), 54.

[42] Yu, J. (2014) Efficient fidelity control by stepwise nucleotide selection in polymerase elongation Abstract: Polymerases select nucleotides. Computational and Mathematical Biophysics, 2(1), 141–160.

[43] Cameron, C. E., Moustafa, I. M., and Arnold, J. J. (2016) Fidelity of Nucleotide Incorporation by the RNA-Dependent RNA Polymerase from Poliovirus. In Enzymes Vol. 39, pp. 293–323 Academic Press.

[44] Huang, J., Brieba, L. G., and Sousa, R. (2000) Misincorporation by wild-type and mutant T7 RNA polymerases: Identification of interactions that reduce misincorporation rates by stabilizing the catalytically incompetent open conformation. Biochemistry, 39(38), 11571–11580.

[45] Arnold, J. J., Smidansky, E. D., Moustafa, I. M., and Cameron, C. E. (2012) Human mitochondrial RNA polymerase: Structure–function, mechanism and inhibition. Biochimica et Biophysica Acta (BBA) - Gene Regulatory Mechanisms, 1819(9-10), 948–960.

[46] Sultana, S., Solotchi, M., Ramachandran, A., Patel, S. S., and Wek, R. C. (2017) Transcriptional fidelities of human mitochondrial POLRMT, yeast mitochondrial Rpo41, and phage T7 single-subunit RNA polymerases. Journal of Biological Chemistry, 292(44), 18145–18160.

[47] Long, C. and Yu, J. (2018) Balancing non-equilibrium driving with nucleotide selectivity at kinetic checkpoints in polymerase fidelity control. Entropy, 20(4), 306.

[48] Romero, M. E., Long, C., La Rocco, D., Keerthi, A. M., Xu, D., and Yu, J. (2021) Probing remdesivir nucleotide analogue insertion to SARS-CoV-2 RNA dependent RNA polymerase in viral replication. Molecular Systems Design and Engineering, 6(11).

[49] Klepeis, J. L., Lindorff-Larsen, K., Dror, R. O., and Shaw, D. E. (2009) Long-timescale molecular dynamics simulations of protein structure and function. Current Opinion in Structural Biology, 19(2), 120–127.

[50] Lazim, R., Suh, D., and Choi, S. (2020) Advances in molecular dynamics simulations and enhanced sampling methods for the study of protein systems. International Journal of Molecular Sciences, 21(17), 1–20.

[51] Long, C., Chao, E., Da, L. T., and Yu, J. (2019) Determining selection free energetics from nucleotide pre-insertion to insertion in viral T7 RNA polymerase transcription fidelity control. Nucleic Acids Research, 47(9), 4721–4735.

[52] Kierzek, R., Burkard, M. E., and Turner, D. H. (1999) Thermodynamics of single mismatches in RNA duplexes. Biochemistry, 38(43), 14214–14223.

[53] Zhang, L., Zhang, D., Wang, X., Yuan, C., Li, Y., Jia, X., Gao, X., Yen, H. L., Cheung, P. P. H., and Huang, X. (2021) 1-Ribose cyano substitution allows Remdesivir to effectively inhibit nucleotide addition and proofreading during SARS-CoV-2 viral RNA replication. Physical Chemistry Chemical Physics, 23(10), 5852–5863.

[54] Abraham, M. J., Murtola, T., Schulz, R., Páall, S., Smith, J. C., Hess, B., and Lindah, E. (2015) Gromacs: High performance molecular simulations through multi-level parallelism from laptops to supercomputers. SoftwareX, **1-2**, 19–25.

[55] Maier, J. A., Martinez, C., Kasavajhala, K., Wickstrom, L., Hauser, K. E., and Simmerling, C. (2015) ff14SB: Improving the Accuracy of Protein Side Chain and Backbone Parameters from ff99SB. Journal of Chemical Theory and Computation, 11(8), 3696– 3713.

[56] Ivani, I., Dans, P. D., Noy, A., Páerez, A., Faustino, I., Hospital, A., Walther, J., Andrio, P., Goñi, R., Balaceanu, A., Portella, G., Battistini, F., Gelpáı, J. L., Gonźalez, C., Vendruscolo, M., Laughton, C. A., Harris, S. A., Case, D. A., and Orozco, M. (2015) Parmbsc1: A refined force field for DNA simulations. Nature Methods, 13(1), 55–58.

[57] Meagher, K. L., Redman, L. T., and Carlson, H. A. (2003) Development of polyphosphate parameters for use with the AMBER force field. Journal of Computational Chemistry, 24(9), 1016–1025.

[58] Bonomi, M., Bussi, G., Camilloni, C., Tribello, G. A., Bańš, P., Barducci, A., Bernetti, M., Bolhuis, P. G., Bottaro, S., Branduardi, D., Capelli, R., Carloni, P., Ceriotti, M., Cesari, A., Chen, H., Chen, W., Colizzi, F., De, S., De La Pierre, M., Donadio, D., Drobot, V., Ensing, B., Ferguson, A. L., Filizola, M., Fraser, J. S., Fu, H., Gasparotto, P., Gervasio, F. L., Giberti, F., Gil-Ley, A., Giorgino, T., Heller, G. T., Hocky, G. M., Iannuzzi, M., Invernizzi, M., Jelfs, K. E., Jussupow, A., Kirilin, E., Laio, A., Limongelli, V., Lindorff-Larsen, K., Löhr, T., Marinelli, F., Martin-Samos, L., Masetti, M., Meyer, R., Michaelides, A., Molteni, C., Morishita, T., Nava, M., Paissoni, C., Papaleo, E., Parrinello, M., Pfaendtner, J., Piaggi, P., Piccini, G. M., Pietropaolo, A., Pietrucci, F., Pipolo, S., Provasi, D., Quigley, D., Raiteri, P., Raniolo, S., Rydzewski, J., Salvalaglio, M., Sosso, G. C., Spiwok, V., Šponer, J., Swenson, D. W., Tiwary, P., Valsson, O., Vendruscolo, M., Voth, G. A., and White, A. (2019) Promoting transparency and reproducibility in enhanced molecular simulations. Nature Methods, 16(8), 670–673.

[59] Price, D. J. and Brooks, C. L. (2004) A modified TIP3P water potential for simulation with Ewald summation. Journal of Chemical Physics, 121(20), 10096–10103.

[60] Essmann, U., Perera, L., Berkowitz, M. L., Darden, T., Lee, H., and Pedersen, L. G. (1995) A smooth particle mesh Ewald method. The Journal of Chemical Physics, 103(19), 8577–8593.

[61] Berendsen, H. J., Postma, J. P., Van Gunsteren, W. F., Dinola, A., and Haak, J. R. (1984) Molecular dynamics with coupling to an external bath. The Journal of Chemical Physics, 81(8), 3684–3690.

[62] Parrinello, M. and Rahman, A. (1981) Polymorphic transitions in single crystals: A new molecular dynamics method. Journal of Applied Physics, 52(12), 7182–7190.

[63] Torrie, G. M. and Valleau, J. P. (1977) Nonphysical sampling distributions in Monte Carlo free-energy estimation: Umbrella sampling. Journal of Computational Physics, 23(2), 187–199.

[64] Kästner, J. (2011) Umbrella sampling. Wiley Interdisciplinary Reviews: Computational Molecular Science, 1(6), 932–942.

[65] Schlitter, J., Engels, M., and Krüger, P. (1994) Targeted molecular dynamics: A new approach for searching pathways of conformational transitions. Journal of Molecular Graphics, 12(2), 84–89.

[66] Kumar, S., Rosenberg, J. M., Bouzida, D., Swendsen, R. H., and Kollman, P. A. (1992) THE weighted histogram analysis method for free-energy calculations on biomolecules. I. The method. Journal of Computational Chemistry, 13(8), 1011–1021.

[67] Grossfield, A. WHAM - Grossfield Lab.

[68] Efron, B. and Tibshirani, R. (1994) An Introduction to the Bootstrap, Chapman and Hall/CRC,.

[69] Michaud-Agrawal, N., Denning, E. J., Woolf, T. B., and Beckstein, O. (2011) MDAnalysis: A toolkit for the analysis of molecular dynamics simulations. Journal of Computational Chemistry, 32(10), 2319–2327.

[70] Waskom, M. L. (2021) seaborn: statistical data visualization. Journal of Open Source Software, 6(60), 3021.

[71] Hunter, J. D. (2007) Matplotlib: A 2D graphics environment. Computing in Science and Engineering, 9(3), 90–95.

[72] Chunhong Long, Ernesto Romero, M., Liqiang Dai, and Jin Yu (2023) Energetic vs. entropic stabilization between a Remdesivir analogue and cognate ATP upon binding and insertion into the active site of SARS-CoV-2 RNA dependent RNA polymerase. Physical Chemistry Chemical Physics, 25(19), 13508–13520.

[73] Poch, O., Sauvaget, I., Delarue, M., and Tordo, N. (1989) Identification of four conserved motifs among the RNA-dependent polymerase encoding elements.. The EMBO Journal, 8(12), 3867–3874.

[74] Shi, J., Perryman, J. M., Yang, X., Liu, X., Musser, D. M., Boehr, A. K., Moustafa, I. M., Arnold, J. J., Cameron, C. E., and Boehr, D. D. (2019) Rational Control of Poliovirus RNA-Dependent RNA Polymerase Fidelity by Modulating Motif-D Loop Conformational Dynamics. Biochemistry, 58(36), 3735–3743.

[75] Alonso, H., Bliznyuk, A. A., and Gready, J. E. (2006) Combining docking and molecular dynamic simulations in drug design. Medicinal Research Reviews, 26(5), 531–568.

[76] Schimunek, J., Seidl, P., Elez, K., Hempel, T., Le, T., Nóe, F., Olsson, S., Raich, L., Winter, R., Gokcan, H., Gusev, F., Gutkin, E. M., Isayev, O., Kurnikova, M. G., Narangoda, C. H., Zubatyuk, R., Bosko, I. P., Furs, K. V., Karpenko, A. D., Kornoushenko, Y. V., Shuldau, M., Yushkevich, A., Benabderrahmane, M. B., Bousquet-Melou, P., Bureau, R., Charton, B., Cirou, B. C., Gil, G., Allen, W. J., Sirimulla, S., Watowich, S., Antonopoulos, N. A., Epitropakis, N. E., Krasoulis, A. K., Pitsikalis, V. P., Theodorakis, S. T., Kozlovskii, I., Maliutin, A., Medvedev, A., Popov, P., Zaretckii, M., Eghbal-zadeh, H., Halmich, C., Hochreiter, S., Mayr, A., Ruch, P., Widrich, M., Berenger, F., Kumar, A., Yamanishi, Y., Zhang, K. Y., Bengio, E., Bengio, Y., Jain, M. J., Korablyov, M., Liu, C.-H., Marcou, G., Glaab, E., Barnsley, K., Iyengar, S. M., Ondrechen, M. J., Haupt, V. J., Kaiser, F., Schroeder, M., Pugliese, L., Albani, S., Athanasiou, C., Beccari, A., Carloni, P., D’Arrigo, G., Gianquinto, E., Goßen, J., Hanke, A., Joseph, B. P., Kokh, D. B., Kovachka, S., Manelfi, C., Mukherjee, G., Muñiz-Chicharro, A., Musiani, F., Nunes-Alves, A., Paiardi, G., Rossetti, G., Sadiq, S. K., Spyrakis, F., Talarico, C., Tsengenes, A., Wade, R. C., Copeland, C., Gaiser, J., Olson, D. R., Roy, A., Venkatraman, V., Wheeler, T. J., Arthanari, H., Blaschitz, K., Cespugli, M., Durmaz, V., Fackeldey, K., Fischer, P. D., Gorgulla, C., Gruber, C., Gruber, K., Hetmann, M., Kinney, J. E., Das, K. M. P., Pandita, S., Singh, A., Steinkellner, G., Tesseyre, G., Wagner, G., Wang, Z.-F., Yust, R. J., Druzhilovskiy, D. S., Filimonov, D. A., Pogodin, P. V., Poroikov, V., Rudik, A. V., Stolbov, L. A., Veselovsky, A. V., Rosa, M. D., Simone, G. D., Gulotta, M. R., Lombino, J., Mekni, N., Perricone, U., Casini, A., Embree, A., Gordon, D. B., Lei, D., Pratt, K., Voigt, C. A., Chen, K.-Y., Jacob, Y., Krischuns, T., Lafaye, P., Zettor, A., Rodŕıguez, M. L., White, K. M., Fearon, D., Delft, F. V., Walsh, M. A., Horvath, D., III, C. L. B., Falsafi, B., Ford, B., Garćıa-Sastre, A., Lee, S. Y., Naffakh, N., Varnek, A., Klambauer, and Hermans, T. M. (2023) A community effort to discover small molecule SARS-CoV-2 inhibitors. bioRxiv,.

[77] Yang, X., Liu, X., Musser, D. M., Moustafa, I. M., Arnold, J. J., Cameron, C. E., and Boehr, D. D. (2017) Triphosphate reorientation of the incoming nucleotide as a fidelity checkpoint in viral RNA-dependent RNA Polymerases. Journal of Biological Chemistry, 292(9), 3810–3826.

[78] Jones, A. N., Mourão, A., Czarna, A., Matsuda, A., Fino, R., Pyrc, K., Sattler, M., and Popowicz, G. M. (2022) Characterization of SARS-CoV-2 replication complex elongation and proofreading activity. Scientific Reports, 12(1).

[79] Petushkov, I., Esyunina, D., and Kulbachinskiy, A. (2023) Effects of natural RNA modifications on the activity of SARS-CoV-2 RNA-dependent RNA polymerase. FEBS Journal, 290(1), 80–92.

[80] Gordon, C. J., Tchesnokov, E. P., Woolner, E., Perry, J. K., Feng, J. Y., Porter, D. P., and Göotte, M. (2020) Remdesivir is a direct-acting antiviral that inhibits RNA-dependent RNA polymerase from severe acute respiratory syndrome coronavirus 2 with high potency. Journal of Biological Chemistry, 295(20), 6785–6797.

[81] Campagnola, G., McDonald, S., Beaucourt, S., Vignuzzi, M., and Peersen, O. B. (2015) Structure-Function Relationships Underlying the Replication Fidelity of Viral RNA-Dependent RNA Polymerases. Journal of Virology, 89(1), 275–286.

[82] Long, C., Chao, E., Da, L. T., and Yu, J. (2019) A Viral T7 RNA Polymerase Ratcheting Along DNA With Fidelity Control. Computational and Structural Biotechnology Journal, 17, 638–644.

[83] Bhakat, S. (2022) Collective variable discovery in the age of machine learning: reality, hype and everything in between. RSC Advances, 12(38), 25010–25024.

[84] Bonati, L., Trizio, E., Rizzi, A., and Parrinello, M. (2023) A unified framework for machine learning collective variables for enhanced sampling simulations: mlcolvar. bioRxiv,.

[85] Wang, J., Arantes, P. R., Bhattarai, A., Hsu, R. V., Pawnikar, S., Huang, Y. M., Palermo, G., and Miao, Y. (2021) Gaussian accelerated molecular dynamics: Principles and applications. WIREs Computational Molecular Science, 11(5).

[86] Chong, L. T., Saglam, A. S., and Zuckerman, D. M. Path-sampling strategies for simulating rare events in biomolecular systems. (2017).

