## Supplemental Information for "Trapping non-cognate nucleotide upon initial binding for replication fidelity control in SARS-CoV-2 RNA dependent RNA polymerase"

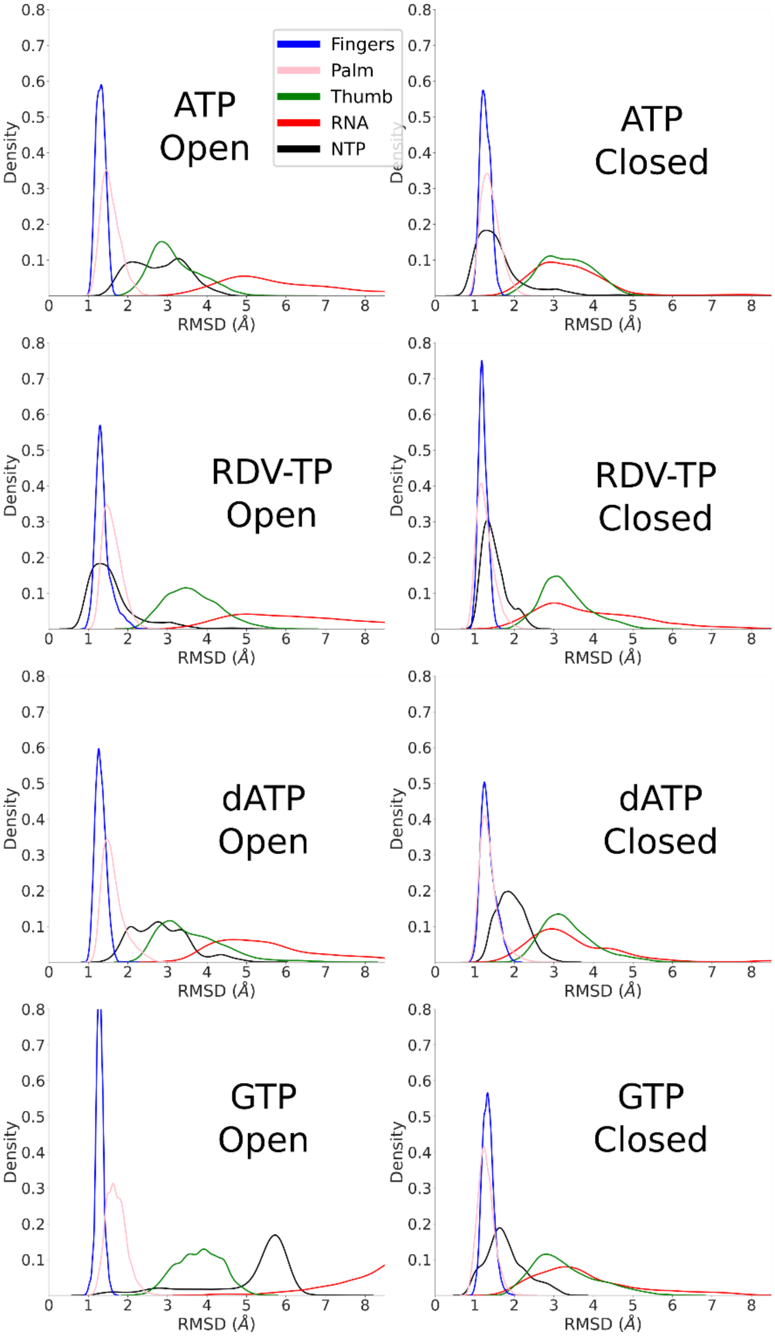


**Figure S1.** The root-mean-square displacements (RMSDs) of SARS-CoV-2 RdRp structural subdomains (backbone atoms), RNA (phosphate backbone), and NTP (heavy atoms) measured from equilibrium ensemble MD simulations (10x100 ns each system). The RMSDs are shown from *top to bottom* for cognate ATP, drug analog RDV-TP, non-cognate dATP and GTP upon initial binding (active site open; *left*) and insertion (active site closed; *right*) states. The subdomains are shown in different colors: fingers (blue). Palm (pink), thumb (green), RNA (red), and NTP (black).

| **Open State** | ATP | RDV-TP | dATP | GTP |
|---|---|---|---|---|
| Motif A | 1.6±0.3 | 1.7±0.5 | 1.7±0.4 | 1.8±0.3 |
| Motif B | 1.3±0.4 | 1.2±0.3 | 1.3±0.3 | 1.2±0.2 |
| Motif C | 1.4±0.5 | 1.3±0.4 | 1.4±0.5 | 0.9±+0.2 |
| Motif D | 1.6±0.4 | 1.6±0.4 | 1.8±0.6 | 1.6±0.3 |
| Motif E | 1.4±0.5 | 1.3±0.3 | 1.6±0.5 | 1.3±0.3 |
| Motif F | 1.6±0.3 | 1.5±0.3 | 1.4±0.2 | 1.6±0.3 |
| Motif G | 1.4±0.3 | 1.3±0.3 | 1.4±0.3 | 1.3±0.2 |

**Table S1.** Average Motif RMSD from the initial binding equilibrium ensemble simulations using the closed state minimized structure as reference. Units are in Angstrom.

| **Closed State** | ATP | RDV-TP | dATP | GTP |
|---|---|---|---|---|
| Motif A | 1.4±0.3 | 1.1±0.2 | 1.3±0.3 | 1.1±0.2 |
| Motif B | 1.0±0.2 | 0.8±0.2 | 1.0±0.3 | 1.1±0.2 |
| Motif C | 0.9±0.2 | 0.8±0.2 | 1.0±0.3 | 0.8±0.2 |
| Motif D | 1.5±0.3 | 1.3±0.3 | 1.4±0.4 | 1.4±0.3 |
| Motif E | 1.2±0.4 | 1.1±0.3 | 1.2±0.4 | 1.0±0.3 |
| Motif F | 1.3±0.2 | 1.1±0.2 | 1.3±0.3 | 1.3±0.2 |
| Motif G | 1.4±0.3 | 1.5±0.3 | 1.6±0.5 | 2.1±0.4 |

**Table S2.** Average Motif RMSD from the insertion equilibrium ensemble simulations using the closed state minimized structure as reference. Units are in Angstrom.

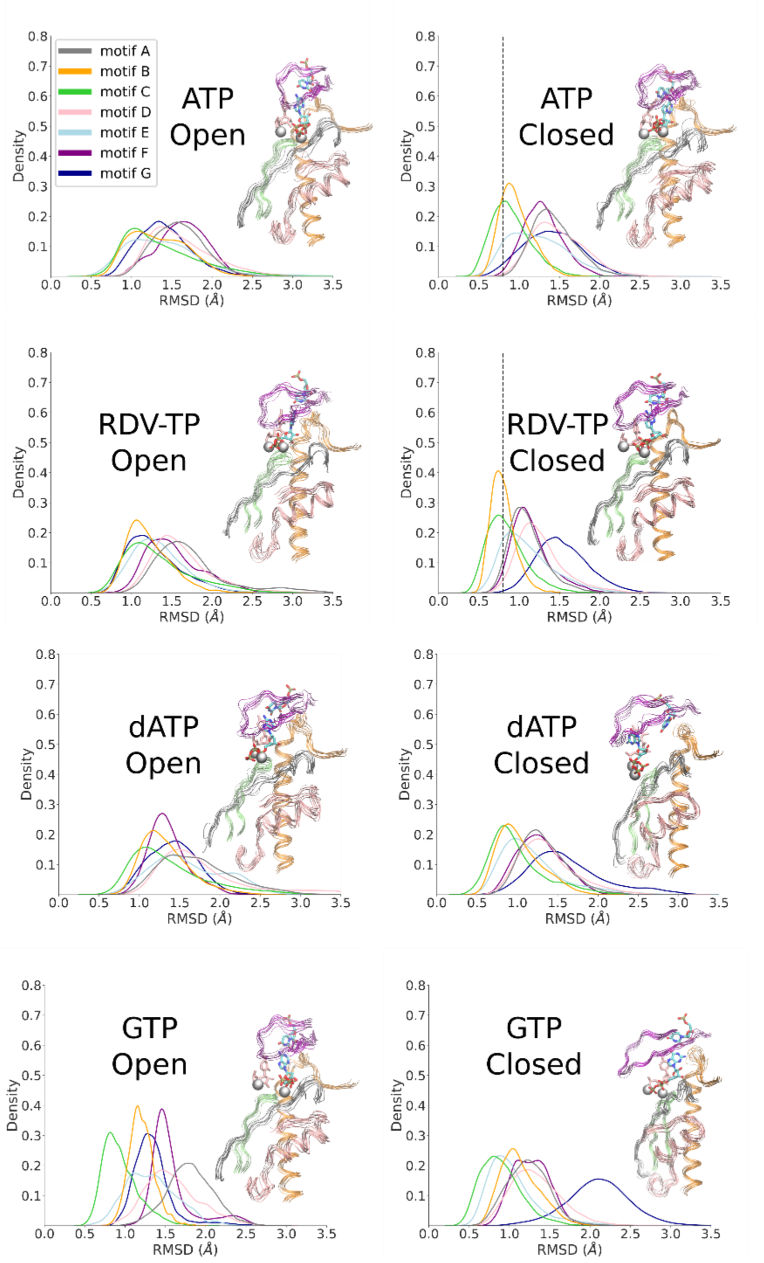


**Figure S2.** The root-mean-square displacements (RMSDs) of RdRp structural motifs (backbone atoms) measured from equilibrium ensemble MD simulations (10x100 ns each system). The motif RMSDs are shown from *top to bottom* for ATP, RDV-TP, dATP, and GTP upon initial binding (active site open; *left*) and insertion (active site closed; *right*) states. The seven key motif RMSD are displayed in different colors: motif A (gray), Motif B (orange), Motif C (green), Motif D (Pink), motif E (light blue), motif F (purple), and motif G (dark blue). Structural representations of those motifs along with NTP, uracil template nucleotide, and two catalytic MG ions are shown for each simulation system. The dotted black line indicates the reference group of motifs (B & C) for systems of inserted ATP/RDV-TP.

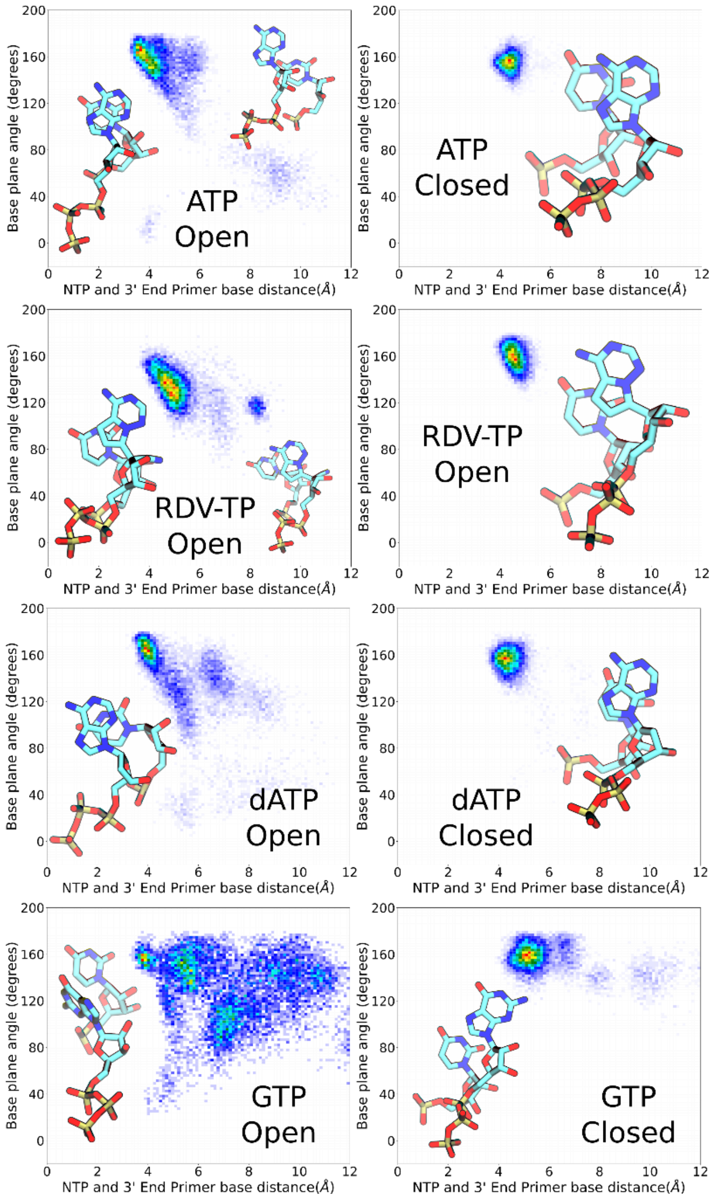


**Figure S3.** NTP and 3’-end primer association geometries sampled from equilibrium ensemble simulations. The geometric measures (see Methods) are shown between the 3’-end primer and individual incoming NTP *from top to bottom*: ATP, RDV-TP, dATP, and GTP are demonstrated, upon initial binding (*left*) and insertion (*right*) for each NTP species. Licorice representations of the NTP and 3’-end primer show the dominant geometries for each simulation system. Distance is measured by the center of mass between the bases. Base plane angle is measured using the C1’-C2-C5 (3’ end primer) and C1’-C7-C5 (NTP).

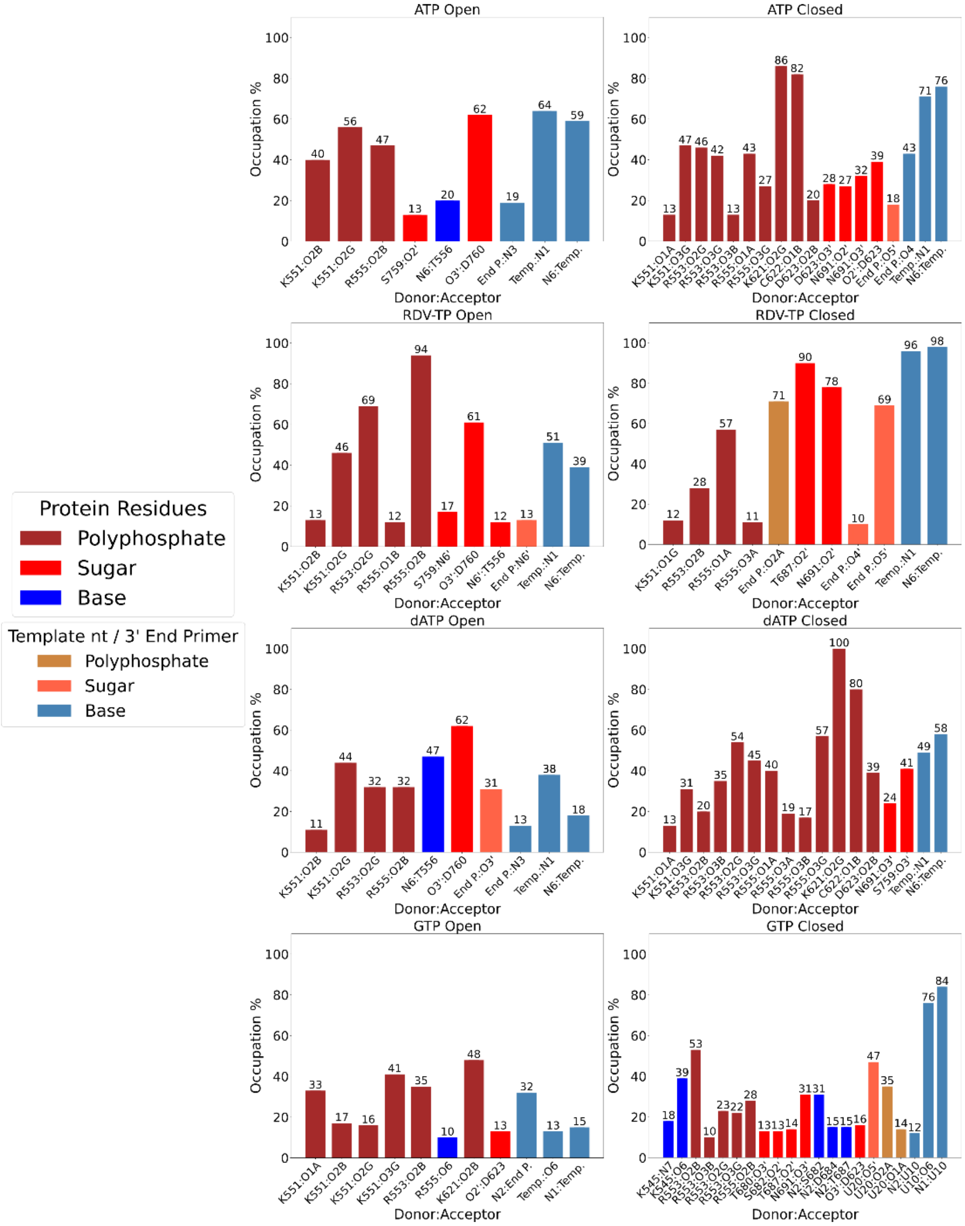


**Figure S4.** The hydrogen bonding (HB) occupancy from the equilibrium ensemble simulations for each NTP, shown from *top to bottom*: ATP, RDV-TP, dATP, and GTP, upon initial binding (open, *left*) to insertion (closed, *right*). Each unique HB interaction >10% population is considered. Two color-code sets are used: protein-NTP interactions use brown (polyphosphate), red (sugar), and blue (base), and for template-nt / 3’ end primer-NTP interactions use light brown (polyphosphate), light red (sugar), and light blue (base).

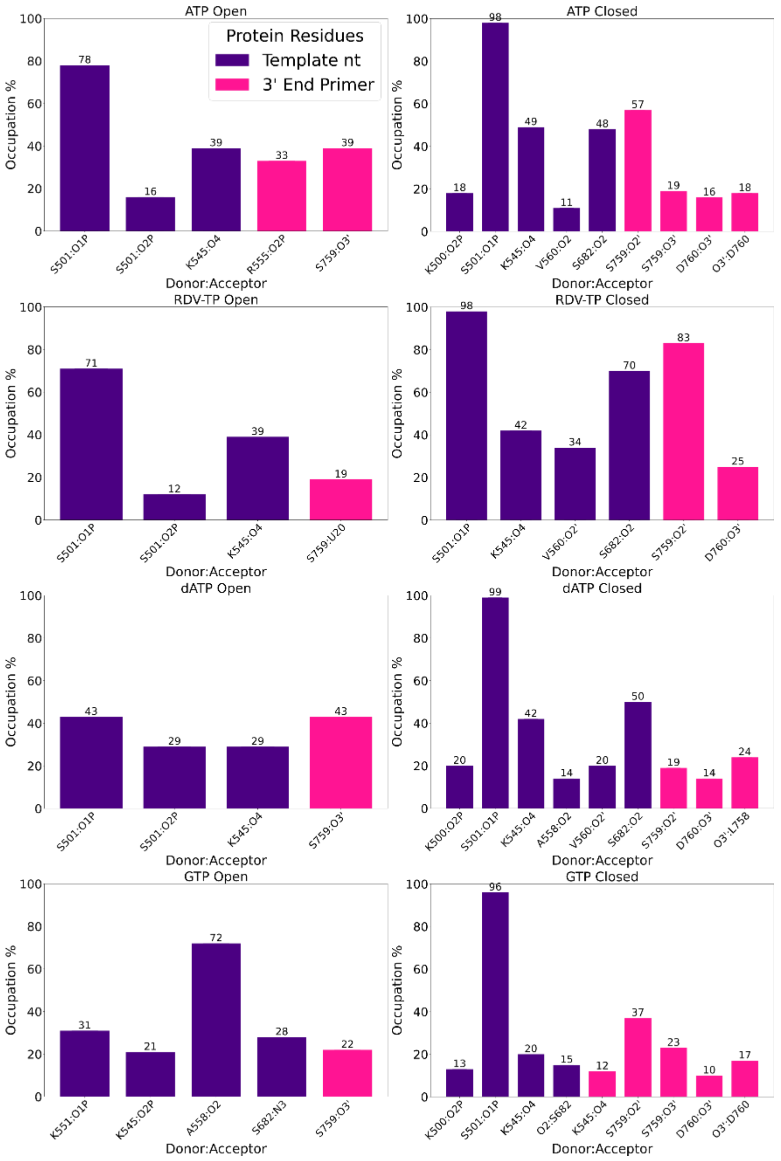


**Figure S5.** The hydrogen bonding (HB) occupancy from the equilibrium ensemble simulations for protein-template nt Uracil (purple) and protein-3’ end primer (pink) for each NTP simulation system, shown from *top to bottom*: ATP, RDV-TP, dATP, and GTP, upon initial binding (open, *left*) to insertion (closed, *right*). Each unique HB interaction >10% population is considered.

**Table S3.** Umbrella Sampling Parameters: force constant $k$ for each path and total number of windows used.

| NTP | Forward $k$ $\left(\frac{kcal}{mol\,Å^2}\right)$ | Backward $k$ $\left(\frac{kcal}{mol\,Å^2}\right)$ | # of windows |
|---|---|---|---|
| GTP | 501 | 501.9 | 24 |
| GTP† | 250 | 501.9 | 24 |
| dATP | 501 | 250.95 | 21 |
| dATP† | 501 | 250.95 | 35 |
| ATP† | 501 | 501 | 26 |
| RDV-TP | 125 | 125 | 21 |

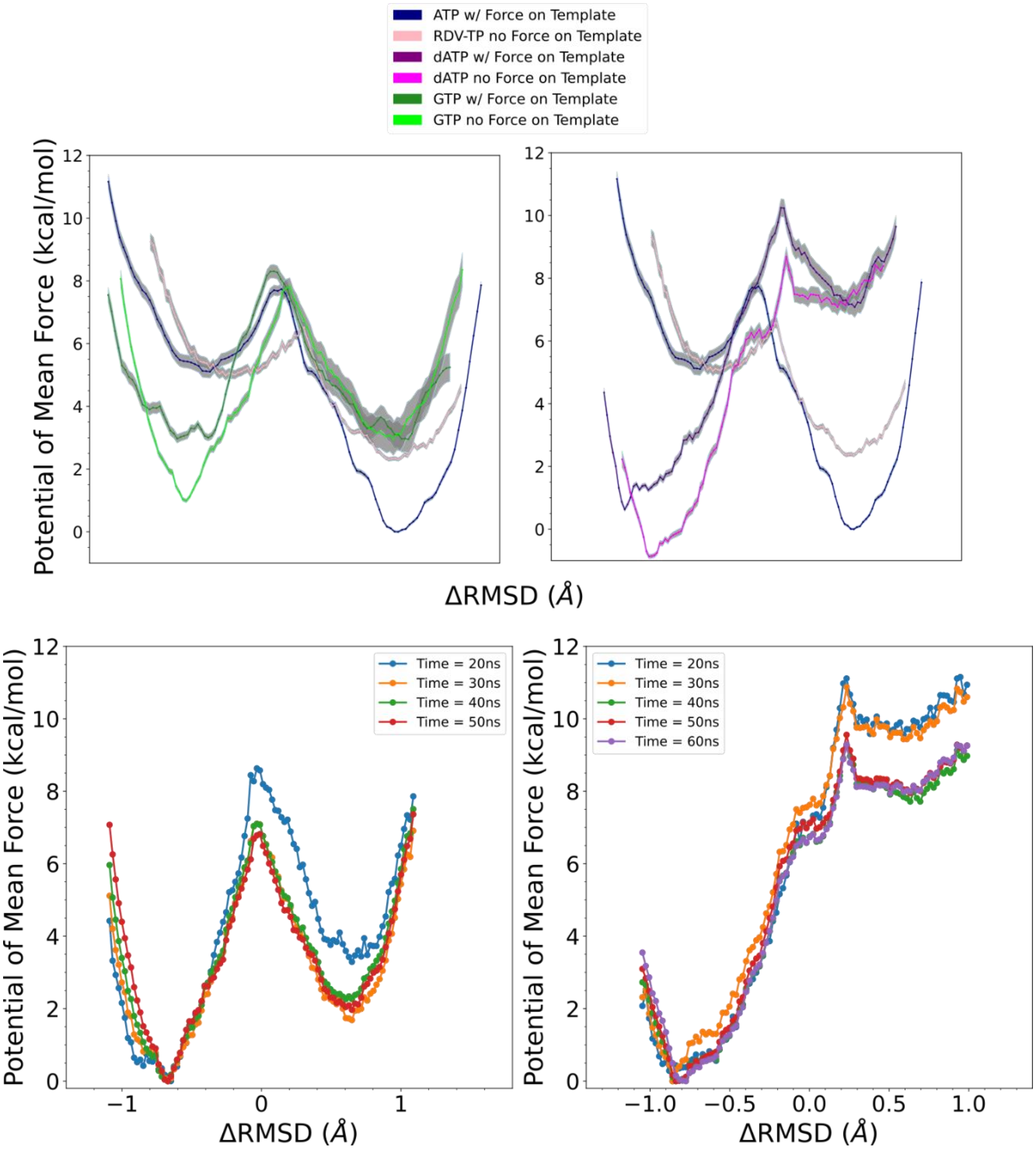


**Figure S6.** The potentials of mean force (PMFs) calculated for various NTPs from initial binding (active site *open*) to the insertion (*closed*) state via umbrella sampling simulations. The difference of RMSDs with respect to *open* and *closed* reference structures $\Delta RMSD \equiv RMSD(X, X_{open})-RMSD(X,X_{closed})$[1], was used as the reaction coordinate in the PMF construction. The *upper left* panel shows the PMFs for GTP, with (dark green) and without (light green) force on the template +1 nucleotide. In both cases, PMFs are shown in comparison with the PMFs obtained for cognate ATP (blue) and drug RDV-TP (pink)[1]. The *upper right* panel shows the PMFs for dATP, with (dark purple) and without (magenta) force on the template +1 nucleotide. The *lower left* and *lower right* panels display convergence plots of PMFs for GTP and dATP, respectively, in current umbrella sampling simulations, without force implemented to the template +1 nucleotide.

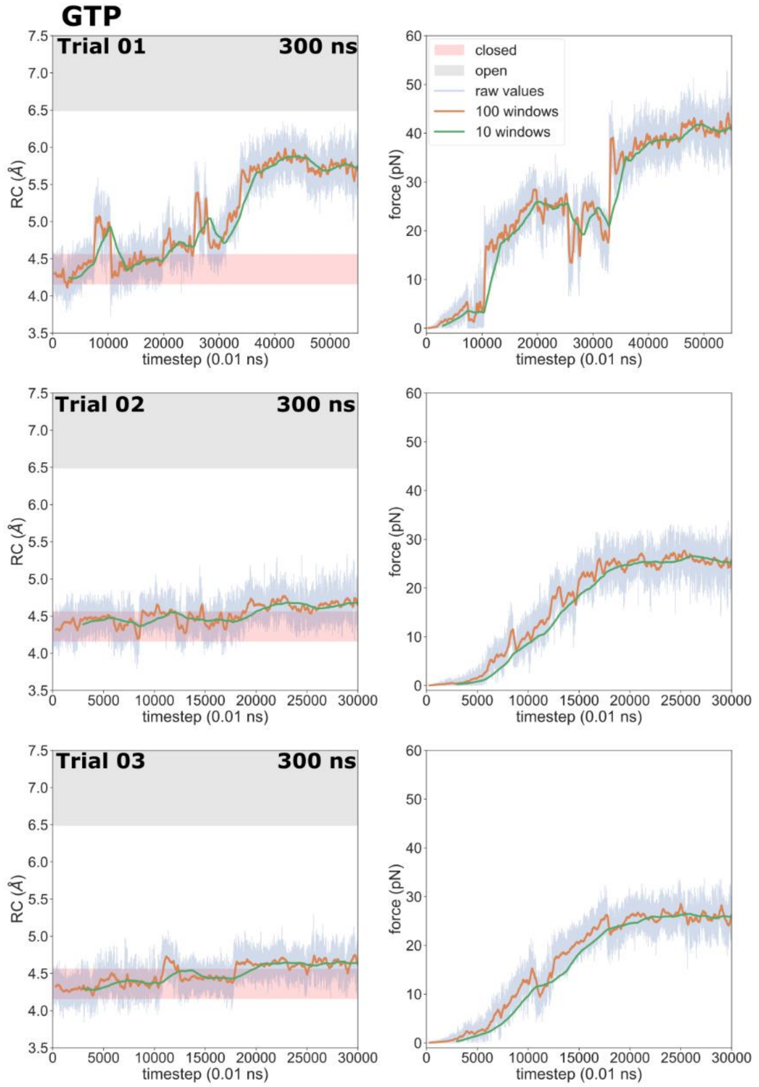


**Figure S7.** Results from steered MD (SMD) pulling GTP from the insertion (active-site closed) state well towards the initial binding (open) state at a rate of 1 Å/ns (force constant 2.4 $\frac{kcal}{mol\,Å^2}$). The reaction coordinate (RC) in pulling simulations, defined as the distance between the GTP and the active center (the center of mass of all αC within 10 Å of the 3' RNA primer), is shown on the left panels. Instantaneous force applied in the SMD simulations is shown on the right panels. Raw data values, 100 window smoothed curves, and 10 window smoothed curves are drawn. ±1 standard deviation of the RCs in the open and closed wells from umbrella sampling are included (as the gray and pink bars on the left panels). Trial simulation 01 was run to a total of 550 ns. The other two trials 02 and 03 were run to a total of 300 ns.

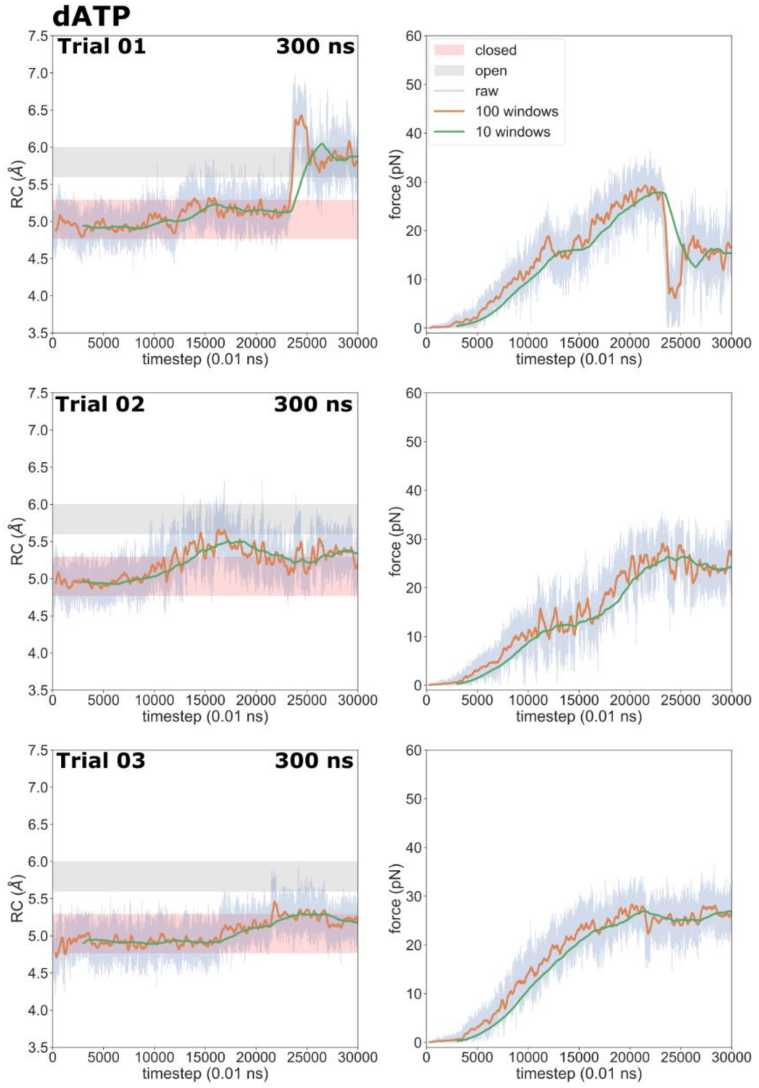


**Figure S8.** Results from steered MD (SMD) pulling dATP from the insertion (active-site closed) state well towards the initial binding (open) state at a rate of 1 Å/ns (force constant 2.4 $\frac{kcal}{mol\,\text{Å}^2}$). The reaction coordinate (RC) in pulling simulations, defined as the distance between the dATP and the active center (the center of mass of all αC within 10 Å of the 3' RNA primer), is shown on the left panels. Instantaneous force applied in the SMD simulations is shown on the right panels. Raw data values, 100 window smoothed curves, and 10 window smoothed curves are drawn. ±1 standard deviation of the RCs in the open and closed wells from umbrella sampling are included (as the gray and pink bars on the left panels). All trial simulations were run to a total of 300 ns.

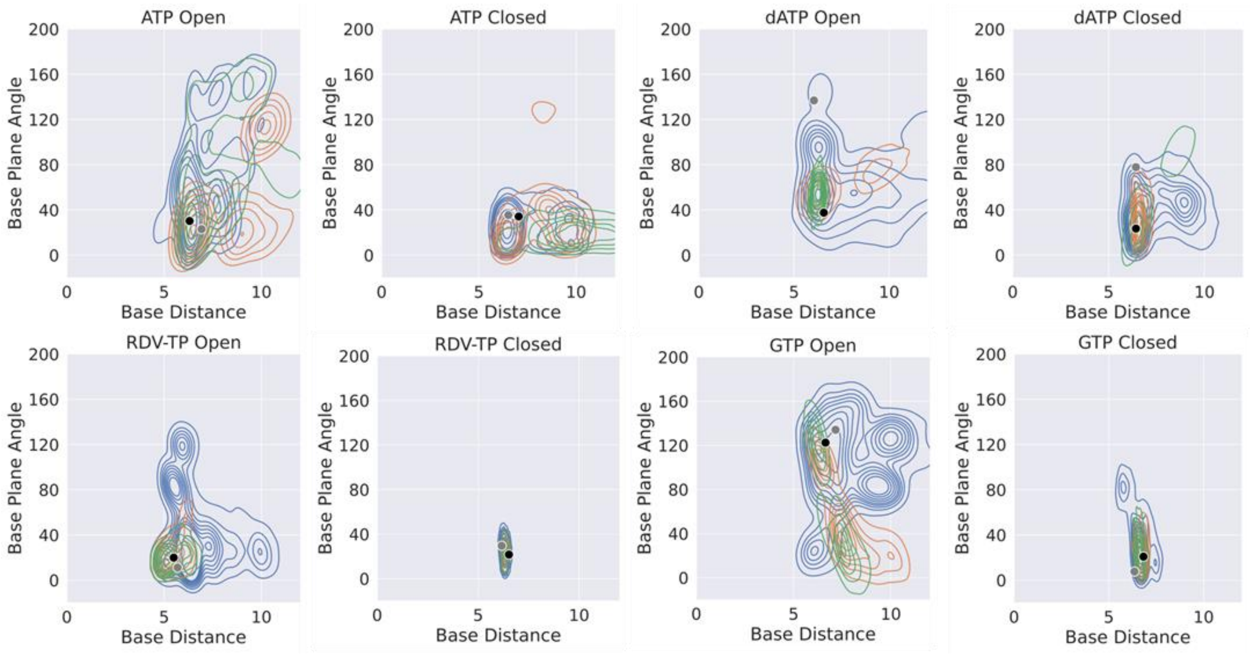


**Figure S9.** The NTP-template association geometry distributions obtained from the umbrella sampling simulations (for PMF calculations) in comparison with that from ensemble equilibrium simulations for various NTP species. A kernel density estimate has been used to visualize the data. Each simulation system (*open* to *closed*, for ATP, RDV-TP, dATP and GTP as in main **Figure 2**), the equilibrium ensemble distribution is shown (blue) along with that obtained from the umbrella sampling (w/ force on template in orange; w/out force on template in green). The black dot indicates the reference state used to generate the initial paths for the umbrella sampling, and grey dot the reference state used in the alchemical calculations[2].

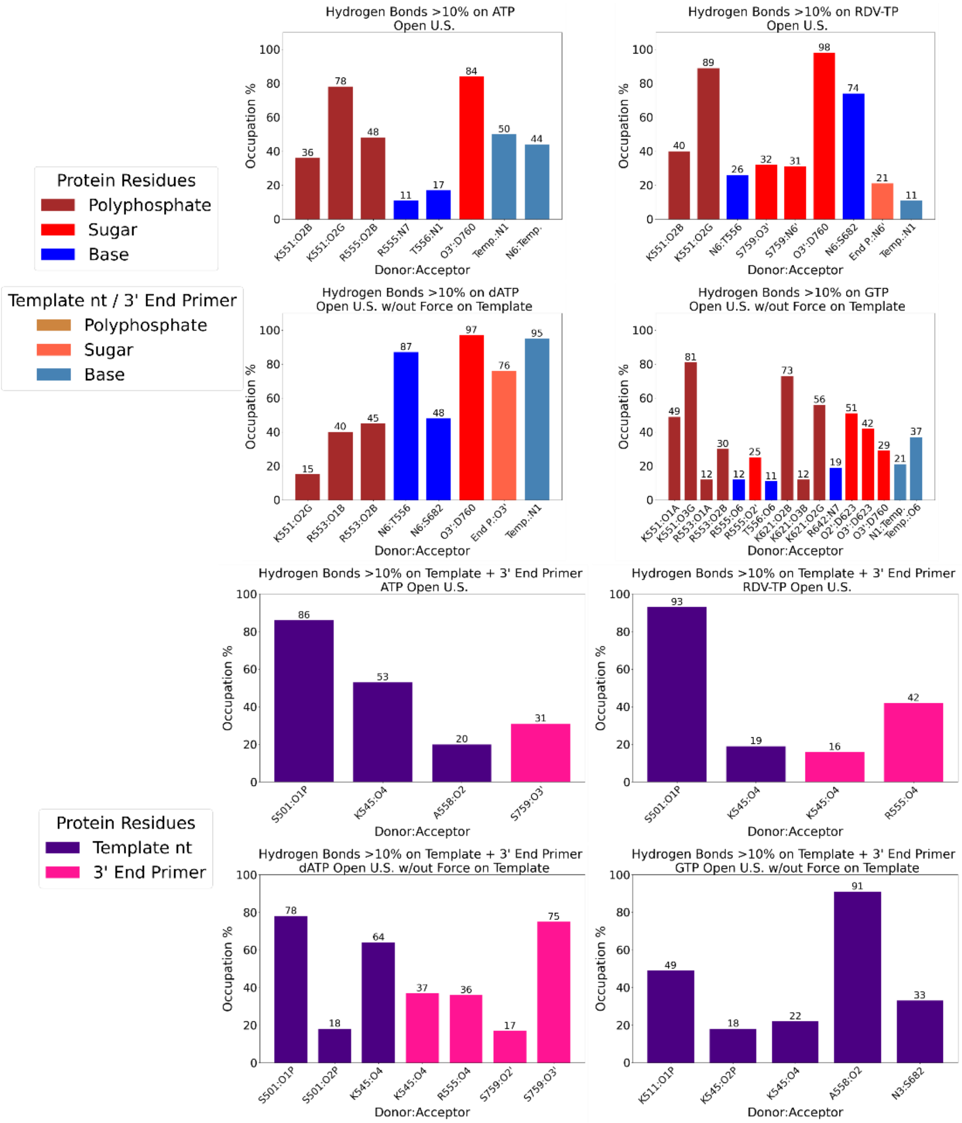


**Figure S10.** The hydrogen bond (HB) occupancy in umbrella sampling trajectories representing the initial binding (open) state minima for each NTP simulation system. The upper panel shows the HBs for each NTP from protein/template +1 nt/3'-end primer. The lower panel shows the protein HBs on the template +1 nt/3'-end primer. For each NTP (ATP, RDV-TP, dATP, and GTP), only unique HB interactions with a population greater than 10% are considered. The same color codes used in Figure S4, and S5 are used to represent the different types of interactions.

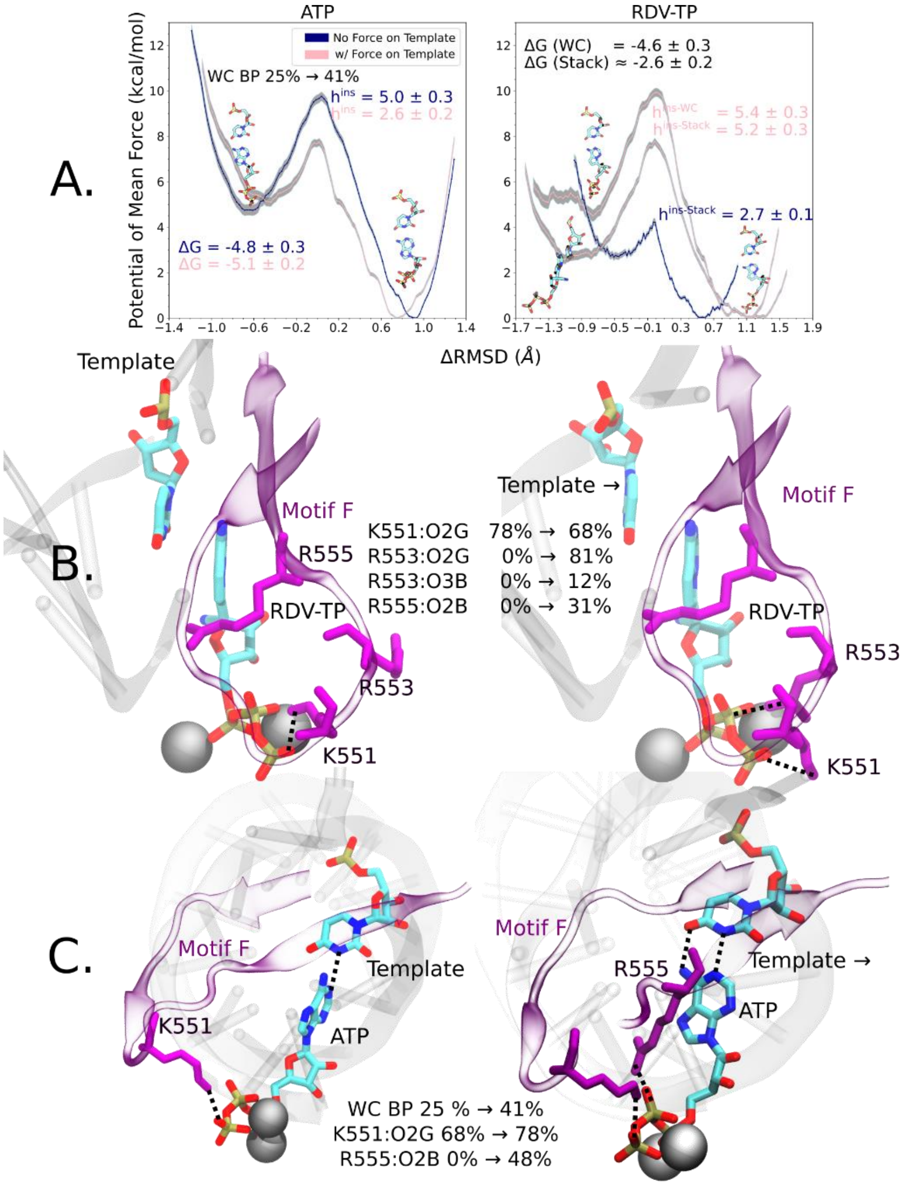


**Figure S11.** A summary of previous work on the insertion PMFs of ATP and RDV-TP[1]. *A:* The PMFs of insertion demonstrate that the free energy barrier ($h^{ins}$) can vary depending on whether there is force implemented on the template +1 nt: w/force, insertion of ATP/RDV-TP results in a lowered/increased barrier comparing with the case w/o force. Labelled energy values are reported in (kcal/mol) *B:* It is shown that starting from the base stacking configuration between RDV-TP and template +1 nt w/o force (*left*), the interactions between the motif F residues (K551, R553, R555) and the polyphosphate are destabilized (i.e., to lower the RDV-TP insertion barrier) comparing to w/force (*right*). *C:* It is shown that implementing the force on the template stabilizes the Watson-Crick base pairing between ATP and the template and enhances the interactions between the motif F residues (K551 and R555) and the polyphosphate, facilitating the insertion (lower the ATP insertion barrier).

1. Romero, M. E. *et al.* Probing remdesivir nucleotide analogue insertion to SARS-CoV-2 RNA dependent RNA polymerase in viral replication. *Mol Syst Des Eng* **6**, (2021).
2. Chunhong Long, Ernesto Romero, M., Liqiang Dai & Jin Yu. Energetic vs. entropic stabilization between a Remdesivir analogue and cognate ATP upon binding and insertion into the active site of SARS-CoV-2 RNA dependent RNA polymerase. *Physical Chemistry Chemical Physics* **25**, 13508–13520 (2023).
